# Uncovering antibody cross-reaction dynamics in influenza A infections

**DOI:** 10.1101/2020.01.06.896274

**Authors:** Gustavo Hernandez-Mejia, Esteban A. Hernandez-Vargas

## Abstract

Uncovering the hidden pathways of how antibodies induced by one influenza strain is effective against another, cross-reaction, is the central dogma for the design of universal flu vaccines. Here, we conceive a stochastic model that successfully represents the antibody cross-reactive data from mice infected with H3N2 influenza strains and further validation with cross-reaction data of H1N1 strains. After modifying several aspects and parameters in the model, our computational simulations highlight that changes in time of infection and the B-cells population are relevant, however, the affinity threshold of B-cells between consecutive infections is a necessary condition for the successful Abs cross-reaction. Our results suggest a reformulation in 3-D of the current antibody influenza landscape.

## Introduction

Seasonal and pandemic influenza A virus (IAV) outbreaks annually cause 3–5 million severe cases and up to 650,000 deaths worldwide (1). Pandemics are commonly caused by viruses with surface glycoproteins to which the human immune system is relatively naive, the haemagglutinin (HA) and neuraminidase (NA) (2), these proteins render antigenic properties of the virus. Antigenically similar viruses circulate for a certain epoch before new strains with modified antigenic characteristics arise and replace them (3, 4). Therefore, the study of antigenic differences is central to uncover insights on the effectiveness of vaccination strategies, the emergence of novel strains, and to understand how immunity induced by one strain is effective against another – the so-called cross-reaction.

The immune response to influenza infection is a set of complex events that include cellular and antibodies (Abs) responses and advanced mechanisms in germinal centers (GCs). GCs are sophisticated compartments were several processes such as B-cell clonal expansion, somatic hypermutation, and affinity-based selection take place (5, 6). Thus, these mechanisms may be crucial for the development of diverse antibody specificities and affinities with potential advantages for broad protection (7–10).

The magnitude and breadth of antibody responses to divergent H3N2 and H1N1 influenza strains due to natural infection in mice, studied by Nachbagauer *et al*. (11), allow to better identify the cross-reaction of Abs. Similar cross-reactivity was observed in memory B-cells between influenza Group 1 (H1) and Group 2 (H3) in humans (12). Importantly, the outcome of Abs cross-reaction responses in (11) may depend on the amino acid (AA) differences of the strains, which follows specific spatial distributions in derived genetic-maps (3, 11, 13). In this direction, extensive studies regarding the Abs cross-reaction within and between influenza strains, acknowledging antigenic differences and data from infections, are needed to guide research in influenza.

Several mathematical models have been proposed to give *in silico* insights of the players in Abs response to infection diseases (14). Most of the models question mechanisms for antibody generation (15) taking into account elements such as B-cells, plasma cells, memory B-cells, the affinity maturation (AM) process in germinal centers, and diverse immunization techniques to promote cross-reactive antibodies (13, 15–18). These models give significant theoretical input of viral and immune properties, however, the integration of experimental data from influenza infection in modeling and challenge remains unexplored.

Here, using a stochastic mathematical model, we study the effects of the naive B-cells repertoire, the antigenic differences, and the time of infection in cross-reaction. While previous studies work in a theoretical shape-space framework, our approach capture experimental data kinetics of specific influenza strains. The model successfully reproduces the general behavior of influenza infection data (11) in terms of breadth and magnitude of antibody response with one or two consecutive infections with H3N2 strains. Importantly, with no modifications in model hypothesis and parameter values, our model can also reflect the cross-reaction outcome of H1N1 strains. We identified that the B-cells undergo affinity modifications that govern Abs cross-reaction, suggesting necessary conditions of affinity rearrangements between infections. Besides, we found that while the antigenic differences between strains play a minor role in consecutive infections, such differences have major effects in simultaneous infections. We further report novel hypotheses regarding representation and modeling of cross-reactive data, accounting for antigenic properties and antibodies distributions.

## Results

We *in-silico* replicate the experimental protocol by Nachbagauer *et al*. (11) for cross-reactive Abs induction in mouse models. The experimental protocol consist of sequential infection with two divergent H1N1 or H3N2 strains of influenza virus. For H1N1 infections, mice were infected with an adapted human seasonal strain A/New Caledonia/20/1999 (NC99), followed by the 2009 human pandemic H1N1 strain A/Netherlands/602/2009 (NL09), six weeks later. NL09 is an isolate antigenically identical to the prototype pandemic H1N1 strain A/California/04/09 (Cal09). The H3N2 infection was first made with a human seasonal strain A/Philippines/2/1982 (Phil82), followed by another human seasonal strain A/Victoria/361/2011 (Vic11), six weeks later. The viral strains in the experiments were chosen to reflect a consecutive exposure history consistent with strains that circulated in humans and because these strains replicate well in mice (11).

The set of computational simulations results are summarized in Fig. 1, which we call the principal test. The test employs 10 simulations for H3N2 (Fig. 1-A-B) or H1N1 (Fig. 1-D-E) influenza strains, separately. The population size of naive B-cells is 10 thousand and the population size of antigen is 10 per strain of infection. As illustrative examples, Abs dynamics of one simulation case with H3N2 strains are depicted in Fig. 1-C and one simulation case with H1N1 strains is shown in Fig. 1-D. Importantly, the H1N1 strains test was made with no scheme modifications from the H3N2 test. The complete set of simulations for H3N2 strains are shown from Fig. (S15) to Fig. (S24). The simulations set for H1N1 strains is depicted from Fig. (S25) to Fig. (S34).

**Fig. 1.**
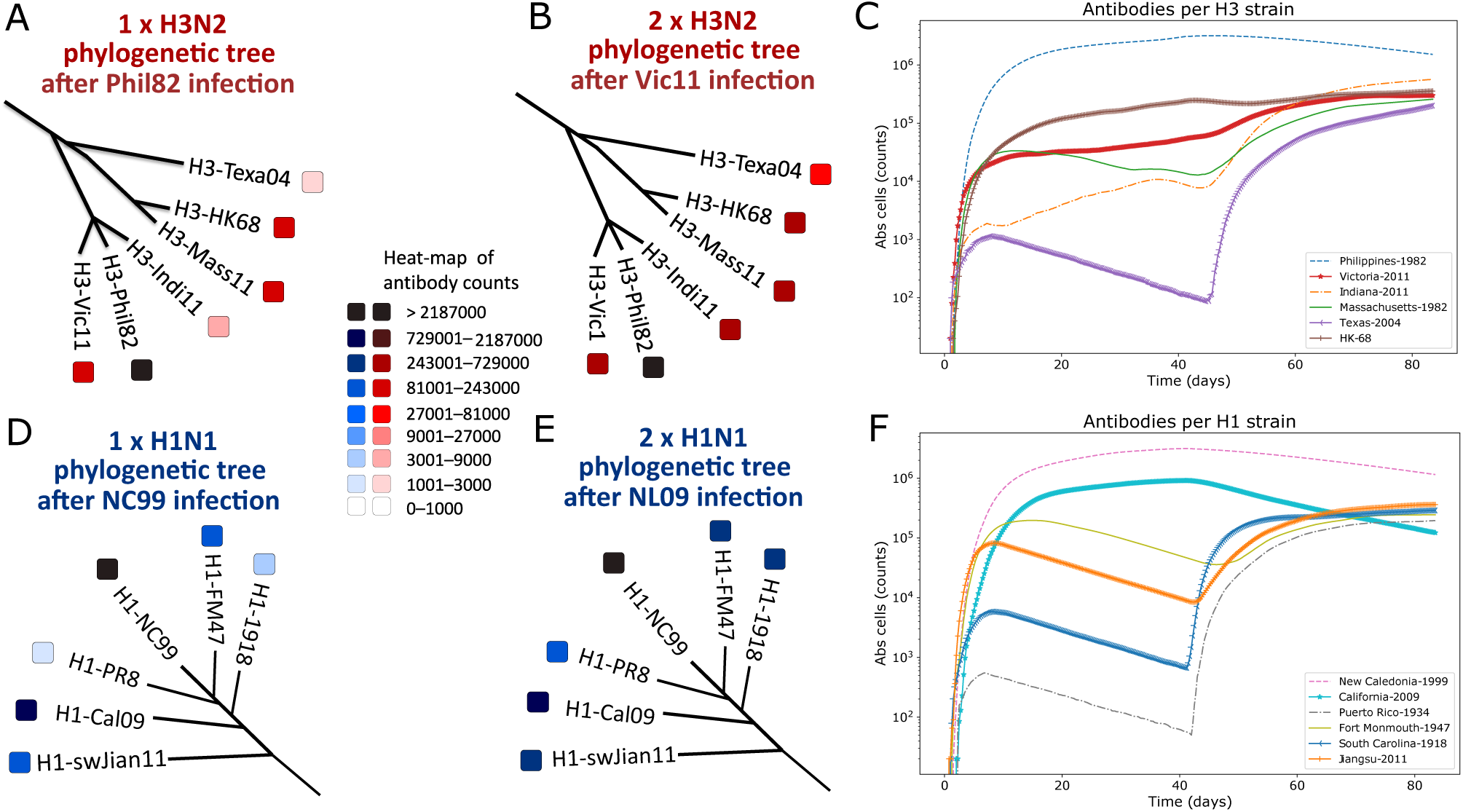
Cross-reactive antibodies response to influenza infections *in-silico,* the principal test. (A-B) A red-colored heat-map display the intervals of Abs counts for different H3N2 influenza strains superimposed onto a phylogenetic tree based on differences in amino acids. (A) Abs counts after the first influenza infection with Philippines-1982 (Phil-82) strain. (B) Abs counts after the second infection with Victoria-2011 (Vic-11) strain. (C) A simulation case of the dynamics of Abs in time (days) for both H3N2 infections, each H3 strain is depicted with different colors and symbols. (D-E) A blue-colored heat-map stand for the intervals of Abs counts for one or two consecutive H1N1 infections whose strains are superimposed onto a phylogenetic tree based on differences in amino acids. (D) The first H1N1 infection is with the New Caledonia-1999 strain (NC99) followed by an infection with the antigenically distinct 2009 pandemic H1N1 (pH1N1) isolate Netherlands-09 (NL09) (E). (F) A simulation case of the dynamics of Abs counts for both H1N1 infections in time (days), each H1N1 strain is depicted with different colors and symbols. The Abs counts represent the result of 10 simulations for H3N2 or H1N1, separately. The corresponding color tone in the squares of each element of the phylogenetic trees stands for the average of Abs counts. The cross-reaction and magnitude of Abs counts represent the ELISA of IgG antibodies to the HA subtypes in serum obtained from experimentally infected mice (11). The time of measurement of Abs counts is consistent with the mice experiments, the Abs response to the first infection is registered 42 days post-infection (dpi) and the results for the second infection are recorded 84 dpi. The phylogenetic tree framework is adapted from Nachbagauer *et al*. (11), the heat-map conserves the scales and color-code. Here, we present Abs as elementary counts, however, the content of phylogenetic trees may be analogous to reciprocal endpoint Abs titers.

### Recapitulating antibodies cross-reactcome

The first H3N2 infection in Fig. 1-A, using the Phil82 strain, promotes the generation of Abs of other H3N2 strains, especially of the Vict11, the A/Hong Kong/1/68 HK68 (HK68) and the A/harbor seal/Massachusetts/1/11 (Mass11) strains. The response of these strains reaches the middle values of the heat-map scale of Abs counts. A more attenuated response can be seen in the A/Indiana/10/11 (Indi11) and the A/canine/Texas/1/04 (Tex04) strains, these strains are within the first three scales of Abs counts. On the other hand, after the second infection with the Vic11 strain, practically all of the H3N2 strains reach high levels of Abs counts, except for the Tex04 strain, as shown in Fig. 1-B. The dynamics of Abs of a simulation case of both H3N2 infections are shown in Fig. 1-C, each strain is associated with a colored-symbol. The fitness of B-cells for antigen increases rapidly during the first GC cycles, following the GCs lose of clonal diversity at disparate rates (26). This behavior is shown in the B-cell dynamics and therefore in Abs of the strain of infection in Fig. 2-B-C, where the matching cells for the Phil82 strain are notably higher compared to the remain strains.

**Fig. 2.**
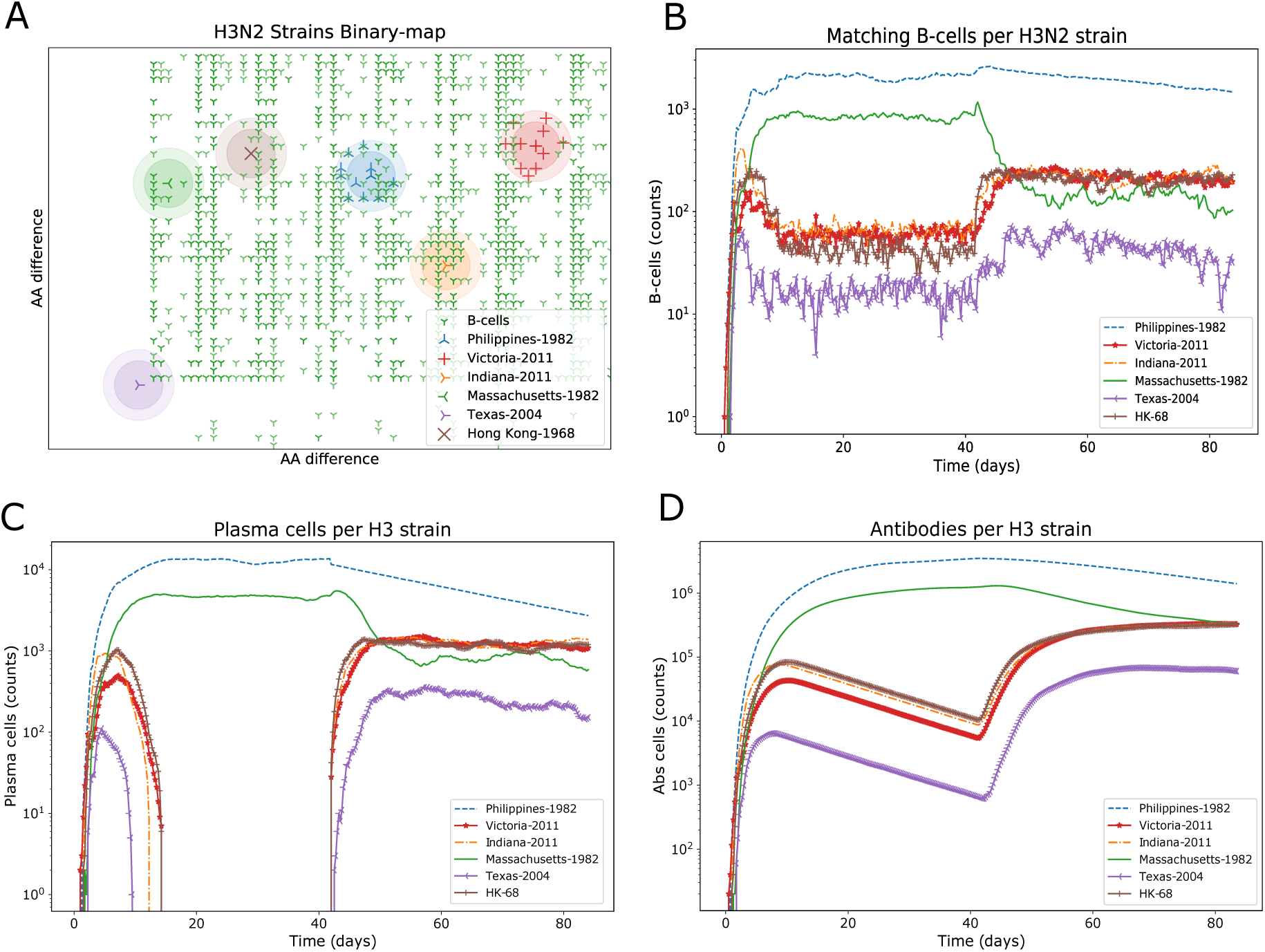
Influenza strains *shape-space* and immune system components dynamics. (A) The genetic binary-map of Fig. 5-A shows now the B-cell population covering most of the H3N2 influenza strains clusters after the Vic11 infection. As a result of AM, the B-cell population targets the H3N2 strains space. In this case, we depict the population shape of B-cells 42 days after the second infection, note that this shape results from the two consecutive infections. (B) A B-cell that falls into the diameter of a strain cluster is considered as a matching B-cell for that strain. The quantity of B-cells inside each cluster is plotted respect to time. (C) A high-affinity B-cell has the chance to generate a plasma cell which decay according to its half-life. Due to a combined effect of the high threshold for the secondary area during the second infection and AM, the plasma cells in the Vic11 infection manage to remain more time than for the Phil82 infection. (D) The antibody population for each strain is generated by plasma cells targeting the corresponding strain. Abs are plotted with respect to time. The four panels (A-D) are the outcome of one of the 10 performed simulations in the principal test.

For the first H1N1 infection in Fig. 1-D using the NC99 strain, Abs cross-reacted mostly with the A/Fort Monmouth/1/47 (FM47), the Cal09 and the A/swine/Jiangsu/40/11 (swJian11) strains. However, an attenuated reaction to the A/South Carolina/1/18 (SC18) and the A/PR/8/34 (PR8) strains was produced. After the second infection with the NL09 strain in Fig. 1-E, all of the strains developed high levels of Abs. A simulation case of the Abs dynamics during both H1N1 infections is shown in Fig. 1-F. The *in-silico* principal test results of Fig. 1 show that the cross-reactive phenomena of natural influenza infection can be captured in both influenza subtypes, maintaining the general shape of breadth and magnitude of Abs response for one or two consecutive infections. These results are comparable to the mice experiments in (11).

### Affinity changes satisfy necessary conditions that direct Abs cross-reaction

The competition between B-cell clones and mutated variants, which include a great number of structural rearrangements (30), is thought to guide the progressive loss of clonal diversity in the B-cell population (29). In this sense, the immune system must regulate the competition to get a balanced level of affinity and diversity. While a rigorous affinity regulation will fastly derive in average population affinity at expenses of diversity, a relaxed regulation will lead to high diversity, but affinity will slowly increase (5, 29). In the model, the cross-reaction outcome of Abs is largely related to the threshold of the common space of the principal and secondary areas, however, there is a greater influence of the threshold of the secondary area (*α*). This reflexes the shape of the binary strings, while the principal area is mostly similar between strains, the secondary area varies according to specific strains. We tested different threshold frameworks for the secondary area to find a mechanism that better represent the affinity–diversity balance of the experimental data of mice.

For both influenza strains, H3N2 and H1N1, we found that a threshold of *α* = 3 during the first infection and a threshold of *α* = 4 for the second infection fairly explain the Abs cross-reactive phenomena in the first and second infections. The threshold of the principal area, *β* = 3, remains unmodified for both infections. Importantly, we found that the threshold modification between infections satisfies conditions of Abs diversity and affinity, which also greatly correlates with the Abs breadth and magnitude the experimental data (11).

Considering a fixed *α* threshold for both infections, we found that using *α* = 3 results in the development of high-affinity strain-specific Abs in the first infection, see results in Fig. (S3). Also, during the second infection, the phylogenetic trees and Abs dynamics of both influenza subtypes inherit a strain-specific behavior from the first infection, which does not correspond with the experimental data. Conversely, using *α* = 4 promotes a wide breadth of the Abs cross-response during the first infection, however, a weak specificity for the strain of infection is present. This behavior is also present in the second infection, inherited from the first one. The outcome of *α* = 4 is depicted in Fig. (S4).

Importantly, the experiments with a fixed threshold guided us to a condition of affinity and diversity balance through the affinity thresholds. We therefore pursuited to satisfy this condition in the model, which resulted in great correlation with the Abs cross-reaction data.

### Antigenic difference weakly affects cross-reaction in consecutive infections with the same influenza A subtype

The strain cluster circles in Fig 5 delimit the affinity for antigen, any B-cell inside a cluster circle is considered to react to the strain of the cluster. The spatial distribution and intersection of influenza clusters may be relevant to the cross-reaction between strains, the Smith approach (13). However, our approach does not consider B-cells in the intersection of clusters as the only ones responsible for cross-reaction since this may derive homogeneous antigen selection. GCs present distinct B-cell clones that lose clonal diversity at diverse rates allowing efficient affinity maturation without homogenizing selection (26). Besides, different modes of viral receptor-binding site recognition arise from diverse germline origins that evolve through diversified AM lines (27). Therefore, the cross-reactive B-cells in our model are those that due to AM can fall inside any cluster circle threshold, ensuring diversification in antigen selection.

Consider the first H3N2 infection with the Phil82 strain whose blue-colored cluster is approximately in the center of the binary-map in Fig 5-A. The closest strains clusters are Vic11, Indi11 and HK68, however, the Abs response to the first infection in Fig. 1-A not only reached these strains but also the Mass11 strain cluster. The largest AA difference concerning the Phil82 cluster is found in the Tex04 strain, this genetic dissimilarity contributes to the limited antibody production of the Tex04 strain. Conversely, the Indi11 strain, which is genetically closer than Tex04, develops slightly higher Abs than Tex04 during the first infection. Besides, the cross-reactive response to the Vic11 strain, located in the right extreme of the H3N2 binary-map, develops high Abs counts to all of the clusters. Interestingly, while for the first infection B-cells mostly cover a narrow y-axis area and a wide x-axis area, for the second infection B-cells equally cover both axes, as shown in Fig. 2-A, a result that weakly relies on the antigenic difference of the strains.

The H1N1 NC99 strain is located in the central upper part of the H1N1 binary-map of Fig 5-B, therefore, according to the antigenic similitude, cross-reactive Abs would equally cover the PR34 and FM47 strains. However, a limited Abs response is developed for the PR34 strain while a high response is generated for the FM47 strain. High antibody responses are also present in the NC99, Cal09, and Jian11 strains, although these last two strains are relatively distanced from NC99 in Fig 5-B. Interestingly, while the more distanced SC18 strain develops a moderated Abs response, the closer PR34 strain has even a slightly minor Abs response. The distance comparison is with respect to NC99 strain. The Cal09 infection produced Abs in practically all of the H1N1 strains including SC18 and Jian11, which are in the left and right extremes of the binary-map in Fig 5-B, respectively. The B-cells activity, the Abs and plasma cells dynamics, and the B-cells map coverage on H1N1 strains can be found in Fig. (S2). The differences in B-cells and Abs responses may follow a hierarchical structure based on different antigenic sites of the HA domains (28), which promote that antigenically distanced strains still arise antibodies and do not fully guide the Abs cross-reactome. Importantly, this phenomenon is captured by the model.

#### Antigenically closed and distanced strains tests

We also tested the antigenic difference effect in the cross-reactive outcome by selecting other H3N2 or H1N1 strains that are genetically closed or distanced each other in the genetic maps in Fig. 5. Antigenically close clusters are called the internal clusters and the distanced clusters are called the external clusters. For the internal H3N2 clusters, we selected the Mass11 strain for the first infection and the HK68 for the second one. The internal clusters in H1N1 are NC99 and FM47 for the first and second infection, respectively. Results of H3N2 are shown in Fig. (S5-A-C,G-H), for H1N1 see Fig. (S5-D-F,I-J). The external clusters for H3N2 infections are Tex04 and Vic11 strains. The external clusters for H1N1 infections are SC18 and Jiang11 strains. Results of external H3N2 clusters are shown in Fig. (S6-A-C,G-H), where a strain-specific shape can be observed. The H1N1 external clusters are shown in Fig. (S6-D-F,I-J). Detailed results can be found in the SI Appendix. The general behavior of the consecutive infections for antigenically closed and distanced strains follows the shape of the principal test, especially during the second infection. The last may add to a weakened effect of the genetic difference in the tested strains.

### Negligible influence of the initial naive B-cell shape repertoire

We target to uncover the impact of the initial naive B-cells population shape. Thus, we tested diverse scenarios of naive B-cells repertoires varying distribution, quantity (minimum 1000 cells), and space coverage. We also explore the newly-generated naive B-cells repertoire per simulation versus a fixed B-cell repertoire employed by several simulations. The overall results may stand for weak consequences of the initial naive B-cells repertoire in the complete cross-reaction scheme for consecutive infections.

First, we test the fixed-shape repertoire generating a set of uniformly randomly produced B-cells. The fixed set is used in a set of ten simulations attempting to show possible differences in cross-reaction outcomes due to a fixed initial set of naive B-cells, see Fig. (S7). Conversely, the newly-generated B-cells framework is actually the principal test. Interestingly, there is not a highlighting difference between fixed-shape in Fig. (S7) and the variable-shape tests of Fig. 1-A-C. Due to the complete bit-map space coverage in both assessments, with equal distribution and logic generation, the B-cell population comparably behaves in both fixed-shape and variable-shape cases. We further tested the naive B-cells population size significance in the Abs cross-reaction by varying the number of B-cells from 1000 cells to 10 thousand cells. The complete panorama of B-cells, PCs and Abs dynamics in Fig. (S8), and the Abs counts in the phylogenetic trees of a naive population with 1000 cells are representative to the landscape of a naive population of 10 thousand cells, proportionally speaking. This test is compared to the principal test.

Finally, we explore diverse random distribution and space coverage of initial naive B-cells. The shape-space of B-cells after the first infection strongly determine the Abs outcome of the second infection. However, the random distribution (uniform and Gaussian) and space coverage of initial naive B-cells weakly influence the complete qualitative picture of cross-reactive antibodies. Fig. (S9-A) shows results of consecutive infection with an initial B-cell repertoire covering the complete bit-space range of Fig. (S1) with uniform distribution. Fig. (S9-B) reports results of a repertoire uniformly distributed but covering only the center of the bit-space. A more limited range is reported in Fig. (S9-C), where the space coverage is limited to the area where the H3N2 strains are located on the map, with uniform distribution. Fig. (S9-D) and Fig. (S9-E) show results of a Gaussian distribution in the center of H3 HAs and the center of the bit-space, respectively.

### Time windows up to six weeks between infections favourably elicit cross-reactive Abs

We explore the Abs cross-reaction outcome for different time windows between infections for both strains applying the second infection 10, 20, and 84 days after the first infection. The Abs measurement time window after the second infection remains in 42 days. After the first infection, we found that the tests with time windows of 10 and 20 dpi present a slightly different outcome concerning the principal test, regarding the Abs outcome from the first infection. For a 10 dpi window, the phylogenetic trees of H3N2 and H1N1 strains register high counts of Abs for the first infection in Fig. (S10-A,D), respectively. This outcome is similar in the window test of 20 dpi in Fig. (S11-A,D) for H3N2 and H1N1, respectively. Of note, we found that the level of Abs after the second infection is comparable for the time windows of 10, 20 and 42 dpi (principal test) in both influenza strains. This results are shown in the phylogenetic trees and Abs dynamics in Fig. (S10-B,C,E,F), Fig. (S11-B,C,E,F) and Fig. 1-B,C,E,F, for the mentioned time windows, respectively.

The test of a time window of 84 dpi for the second infection in Fig. (S12) shows a limited Abs outcome for the first infection in H3N2 and H1N1 strains. However, during the second infection, the Abs arise at medium to high levels with respect to the principal test results.

### *Commutative* test

We tested the Abs outcome employing the same H1N1 or H3N2 strains but in the opposite infection order respect to the principal test. In the case of H3N2 strains, the first infection in this test is with Vic11 and the second infection is with Phil82. For H1N1, the first infection is with the Cal09 strain followed by the NC99 strain. The time window between infection remains in 42 d. We found that the Abs outcome in the first infection produces a slightly lower magnitude of cross-reactive Abs, however, as can be observed in Fig. (S13-A,D), the breadth of cross-reaction covers almost all of the H3 or H1 strains, respectively. After the second infection, the general Abs outcome observed in Fig. (S13-B,E) is comparable to the principal test. Summarizing, for the H1N1 and H3N2 strains employed in this and the principal tests, a *commutative* outcome can be observed between these influenza strains.

### Cross-reactive Abs in simultaneous infections

Contrasting outcome scenarios have been questioned for HIV infection, while broad protection can be limited when employing simultaneous infections with different antigen strains (17), the competitive exclusion of broad Abs may be reduced in infections initialized with multiple strains (15). Therefore, we evaluate a simultaneous scheme of influenza infection using the H3N2 or H1N1 strains from the principal test. Importantly, results in Fig. (S14) show that, contrasting with the consecutive infections of Fig. 1, the AA difference may influence closely related H3N2 strains (Vic11, Phil82, Mass11, and HK68, sharing the y-axis) to effectively cross-react with the simultaneous infection strains (Vic11 and Phil82). Besides, distanced strains in H1N1 infections (NC99 and Cal09, sharing the x-axis) results to follow a strain-specific shape, with low to medium Abs count of the remain H1N1 strains. Results suggest that the antigenic similitude between antigens may influence cross-reaction when using a scheme of simultaneous infections but not when applying a consecutive strategy.

## Discussion

The *in silico* capacity to reproduce the mice experiments (11) gives testimony of the competency of the model regarding the direct analogy with Abs cross-reaction and the indirect study of the stochastic nature of GCs, AM of B-cells, and affinity adaptation properties. The model depicts the Abs response to infections with two of six H3N2 strains. Without modeling modifications, the challenge of the model to reproduce the response of H1N1 strains cross-reaction is also favourable, highlighting the model reliability The only difference between H3N2 and H1N1 strains experiments is the introduction of different coordinates in the genetic maps and naturally the change of antigen of infection. Taking these properties, we consider that the model can serve as a tool for future research in infectious diseases.

The mice experiments show that the magnitude of Abs in the HA head domain is higher than those of the stalk domain, with both domain Abs increasing in the second infection. The immunodominance of the HA head domain over the stalk domain, with preferred antigenic sites, may explain these differences (2, 28). A proposed mechanism suggests restricted accessibility of the membrane-proximal HA stalk domain to membrane-bound B-cell receptors, however, it is still not clear what governs these phenomena (31). The immunodominance effect is present especially from the first to the second infection in the mice experiments (11). In this sense, one of the most important questions was how to set the model to enhance the Abs cross-reaction from the first to the second infection. The affinity threshold change in the secondary area between infections boosts cross-reactive Abs during the second infection and allows an effective response to the first infection, aided by a fixed threshold in the principal area. Although the conception of our study does not consider differences between HA domains in modeling, after analyzing results, the model suggests that the secondary area may be analog to the head HA domain and the principal area to the stalk domain. The affinity change may be then related to immunodominance preferences between HA head and stalk domains. Of note, as the HA stalk domain can undergo limited drift under immune pressure (32) and aided by a fast-evolving of the HA head domain (33), the model must incorporate such evolving properties to understand how cross-reaction is modified between infections and the related forces of HA domains in different time scales.

Smith *et al*. (13) proposed that simultaneous infection with distanced (external) antigens considers only B-cells inside the cluster circle to develop Abs for the antigen of infection as shown in Fig 3-A. In this fashion, if the second antigen is sufficiently distanced from the first antigen, there will be a poor B-cell reaction for the second antigen. For an effective cross-reaction of both antigens, the intersection of the clusters circles of both antigens is necessary. This may be more effective with closely related (internal) antigens. In both cases, internal and external clusters, the antigenic distance between antigens plays an important role in cross-reaction. On the other hand, in Fig 3-B our approach does not consider B-cells in the intersection of strain clusters as the only responsible for cross-reaction since this may generate a homogeneous B-cells reaction. The cross-reactive B-cells in our model are those that are inside any strain cluster affinity threshold. The antigenic difference is also important in this approach, however, a more heterogeneous B-cells reaction will lead to better cross-reaction held by data. The proposed cross-reaction framework suggests a correlation with recent studies supporting that GC reactions to complex antigens, such as influenza, permit a range of specificities and affinities, with potential advantages for broad protection (7).

**Fig. 3.**
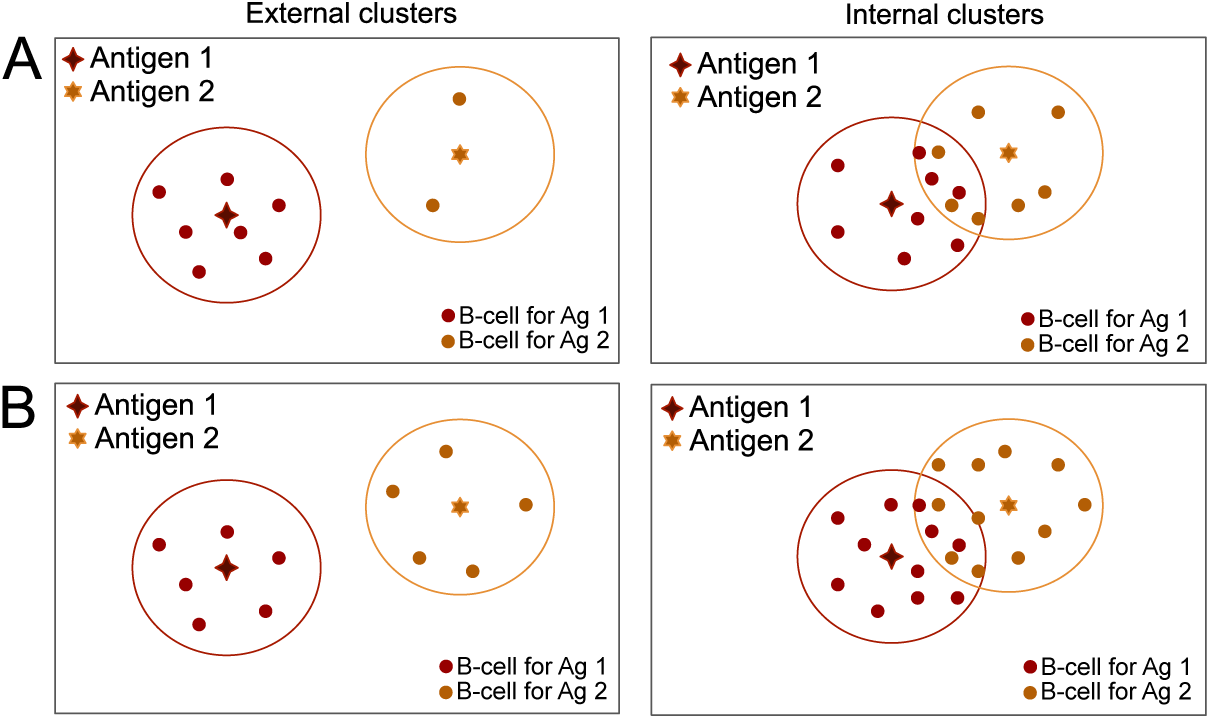
A caricature of the cross-reaction threshold differences between Smith *et al*. (13) and our approach for internal and external clusters. An infection with Antigen 1 develops cross-reactive B-cell responses with Antigen 2 depending on the similitude of antigens and its cluster affinity thresholds. (A) In the Smith approach, the distanced antigens clusters, external clusters, an infection with Antigen 1 develops a B-cell response mostly for itself. In the internal clusters, as the affinity circles of Antigen 1 and 2 are closely related, the cross-reactive B-cells are those in the intersection of both cluster affinity thresholds. (B) In the framework herein presented, an infection with Antigen 1 will develop B-cell responses for itself and also for the distanced Antigen 2, although with minor magnitude. An infection with Antigen 1 in internal clusters will provoke a reaction of B-cell for both antigens with slight differences in magnitude. In both cases, the B-cells that are inside each cluster affinity threshold are considered to be a result of the antigenic difference and the stochasticity in B-cells response.

Further informative results rely on the time windows between infections, the commutation of strains and the naive repertoire of B-cells. A six weeks maximum time window between infections resulted to favor Abs development and better cross-reactive outcome, compared to a twelve weeks window. Also, the Abs response is active within the first 10 h after infection in all experiments. This point suggests a correlation with theoretical studies in CGs function, where high-affinity antibodies appear one day earlier and the PCs population is considerably large (24). On the other hand, regarding the commutive outcome of infections, the static setting of the viral strains may aid in the shape of the commutative results; differences in antigen mutation rate may affect the commutative outcome. Influenza mutation rate varies from 7.1×10^−6^ to 4.5×10^−5^substitutions per nucleotide per cell infection cycle (34) and although reassortment between lineages of the same subtype, the so-called intrasubtype reassortment, is limited by genetic compatibilities of the viruses (35), these changes may handle the commutative shape with, for instance, reordering among H3N2 viruses (36). A comparison of different influenza strains mutation rates during the time scales of infection is further needed to allow us to speculate on the commutation results. Concerning the initial naive B-cells repertoire, although different shapes were tested, we still speculate the influence of other unmodeled forces that may change the cross-reactive panorama making it dependent on the initial repertoire. Recent studies indicate a strong influence of the first infections early in the life of an individual to guide immune responses of later infections in life (37). Our results suggest a comparable behavior since the first infection outcome strongly affects the complete Abs breadth and magnitude of the second infection and, for the tested period of time, the shape is inherited till the end of the simulations.

Recent research in ferrets and humans show that cross-reactive Abs are developed not only within influenza groups but also between them (11, 12). Since our modeling framework shows the cross-reaction in strains of either influenza group 1 (H1N1) or group 2 (H3N2), we consider that the spatial modeling of strains in genetic maps should be improved acknowledging a third dimension in the map. Ito et al. suggest that influenza viruses have followed a gnarled-like evolutionary pathway with an approximately constant curvature (38), showing three-dimensional pathways for H3 or H1 strains, separately. However, this pathway scheme may be unclear in the 2-D representation of genetic maps in Fig. 5. Therefore, we contemplate that similar 3-D shapes of strains positioning of both influenza groups, all together in one three-dimensional map, should be more accurate to stand for cross-reaction between influenza groups. A hypothesized-like map is shown in Fig. 4, where a heat-colored surface cover both influenza groups strains and the AA differences are mapped into three axes. The breadth and magnitude of Abs cross-reaction between and within groups are represented by the surface where the colors stand for the Abs magnitude of each strain and the surface itself represent the breadth of Abs outcome. The dark-blue in the surface of Fig. 4 may highlight strain-specific Abs while the light-blue stands for cross-reactive Abs. The surface is fitted to the AA differences of all strains in the 3-D map. This approach may add to the conception of antibody landscapes, where the x- and y-axes stand for antigenic differences and the z-axis depicts the Abs response (39), however, the surface approach incorporates insights of interactions of influenza groups and cross-reaction of possibly all strains, as well as limitations for it. We hypothesize that the 3-D cross-reactive surface technique will give detail to key questions such as the scope of Abs breadth between groups, the variability due to antigen mutations, specific forces that vary in different animal and human models influencing species-specific immunodominance (40), and hierarchical changes of the immune response to sites of HA (28).

**Fig. 4.**
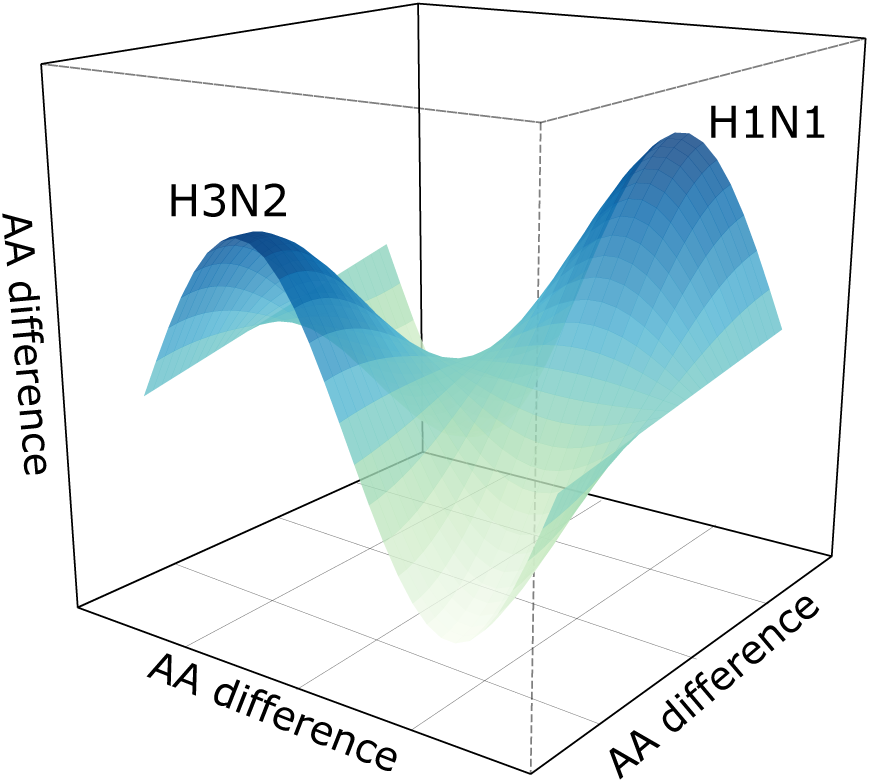
The hypothesized Abs cross-reaction surface. The breadth and magnitude of Abs cross-reaction between and within groups are represented by the 3-D surface where the colors stand for the Abs magnitude of each strain and the surface itself represent the breadth of Abs outcome. The dark-blue in the surface may highlight strain-specific Abs while the light-blue stands for cross-reactive Abs. The surface is fitted to the AA differences of all strains in the 3-D map considering influenza group 1 (H1N1) and group 2 (H3N2).

## STAR Methods

The model herein developed follows the line of previous studies, the philosophy of AM and B-cell fitness is founded in Luo *et al*. (15) and Khailaie *et al*. (18). The principles of antigenic/genetic distance and the shape representation of B-cells and pathogens follow the studies of Smith *et al*. (3, 13) and Lapedes *et al*. (19). The different stages of cell populations going from naive B-cells to Abs is founded in Chaudhury *et al*. (16).

### A. Genetic binary-map

Using the data of experiments with infected mice and the genetic maps in (11), we use the corresponding x- and y-coordinates of each influenza strain to create the genetic binary-maps in Fig. 5. We define a binary metric for the strain’s x- and y-coordinates using strings of binary characters. A binary string of size 2*n* represents the x- and y-coordinates of an antigen or B-cell, therefore, each coordinate is given by a binary string of size *n*. The binary string representation of the coordinates of an antigen and B-cell are shown in Fig. 6. The genetic binary-maps of Fig. 5 shows the spatial localization of each H3N2 or H1N1 influenza strains regarding its AA differences. The affinity-space of a strain is delimited by color-coded clusters (circles) surrounding the strain centroid, the diameter of a cluster is 10 units of AA difference. This diameter is reasonably consistent with the cluster transition average of 13.2 units of AA difference between H3N2 strains (3).

**Fig. 5.**
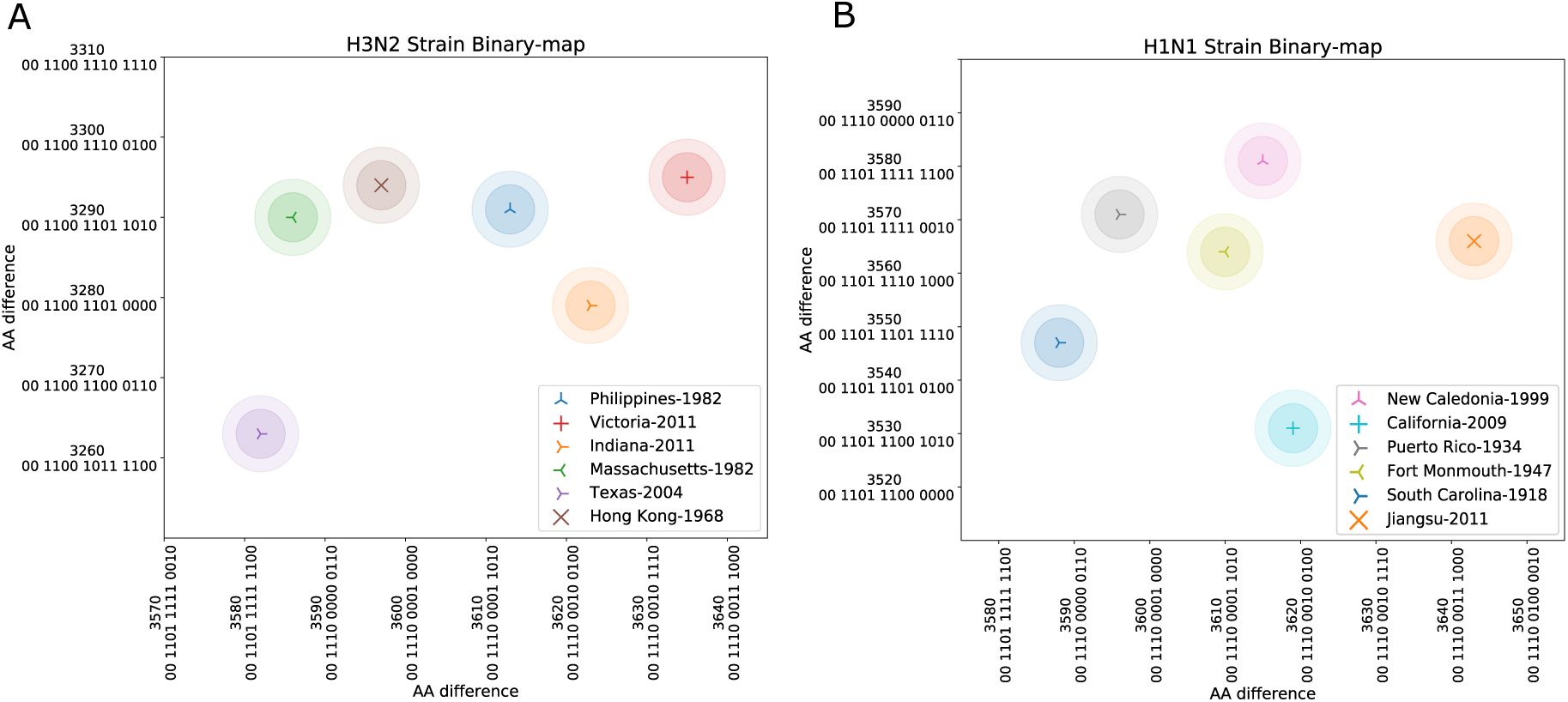
Genetic binary-map. The decimal numbers (base-ten) are accompanied by the binary-system numbers in both axes of the genetic binary-maps. A) Different color-coded symbols stand for the centroid of each H3N2 influenza strains, the centroid is surrounded by the strain cluster coded with the same color. The circle of each strain cluster delimits the affinity-space of the strain. The x- and y-axes show the genetic distance between strains given by the amino acid (AA) difference. B) Diverse color-coded symbols stand for the centroid of each H1N1 influenza strains and a circle represents the cluster of the strain. The genetic maps in this figure are adapted from the antibody landscapes in (11).

**Fig. 6.**
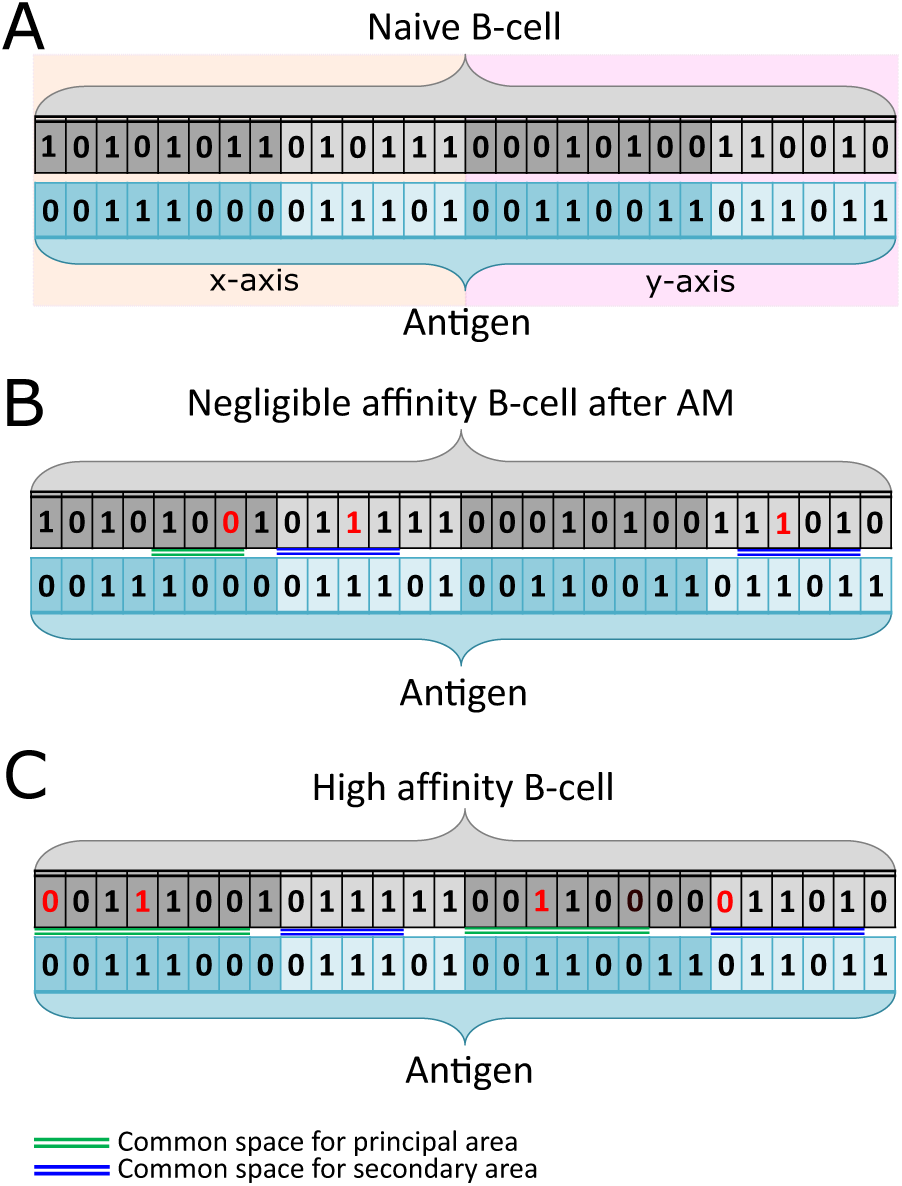
Mathematical model of the evolving affinity of B-cells for antigen. Antigens and B-cells are formed by two strings of 14 binary characters, each string stands for the x- and y-coordinates from the genetic binary-maps. A Binary string is formed by the principal area, the darker one, and the secondary area with a lighter color. The affinity of a B-cell for antigen considers the common space in the principal (green) and secondary (blue) areas. An outstanding affinity considers the longest consecutive common bit-alignments between the antigen and B-cell. A) The naive B-cell population has a very low or no affinity for the antigen. B) Due to the affinity maturation (AM) process, a new set of slightly higher affinity B-cells arise, which establish a new threshold of affinity for next generations of B-cells. Red characters result from several iterations of AM. C) As AM goes on, B-cells reach high affinity for the antigen, characterized by longer common spaces of the principal and secondary areas. The complete scheme of the immune system model is depicted in Fig. 7.

We converted the coordinates values of Fig. 5 from the decimal numeral system (base-ten) to binary values followings the two’s complement approach, a method for representing signed (positive and negative) values in a binary numbering system (SI Appendix1). Using two’s complement, the axes of the binary-map (bit-space in Fig. (S1)) cover a range of 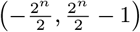, where 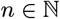 is the binary string size of the axes and works as a range-tuning parameter. In this fashion, the binary strings combinations representing influenza strains naturally holds information regarding the AA differences between strains.

### B. Germinal Center model

We aim to explore the mechanisms of antibody generation that operate in both directions, high fitness to the given antigen and Abs diversity for broad protection, reflecting the AM process that B-cells undergo in GCs (5, 15, 20). The schematic of the AM process is shown in Fig. 6, depicting the antigen and B-cell binding interface with the principal stages of the phenomena we want to investigate. A complete scheme of our model, involving the considered immune elements, is depicted in Fig. 7. While the antigen binary-strings directly come from the strain’s coordinates in the genetic binary-map, the characters of the binary strings representing the naive B-cells population are generated uniformly at random. This approach results in a population of naive B-cell whose binary-strings cover the complete bit-space of Fig. (S1) and can be mapped in a genetic binary-maps as the ones of Fig. 5.

**Fig. 7.**
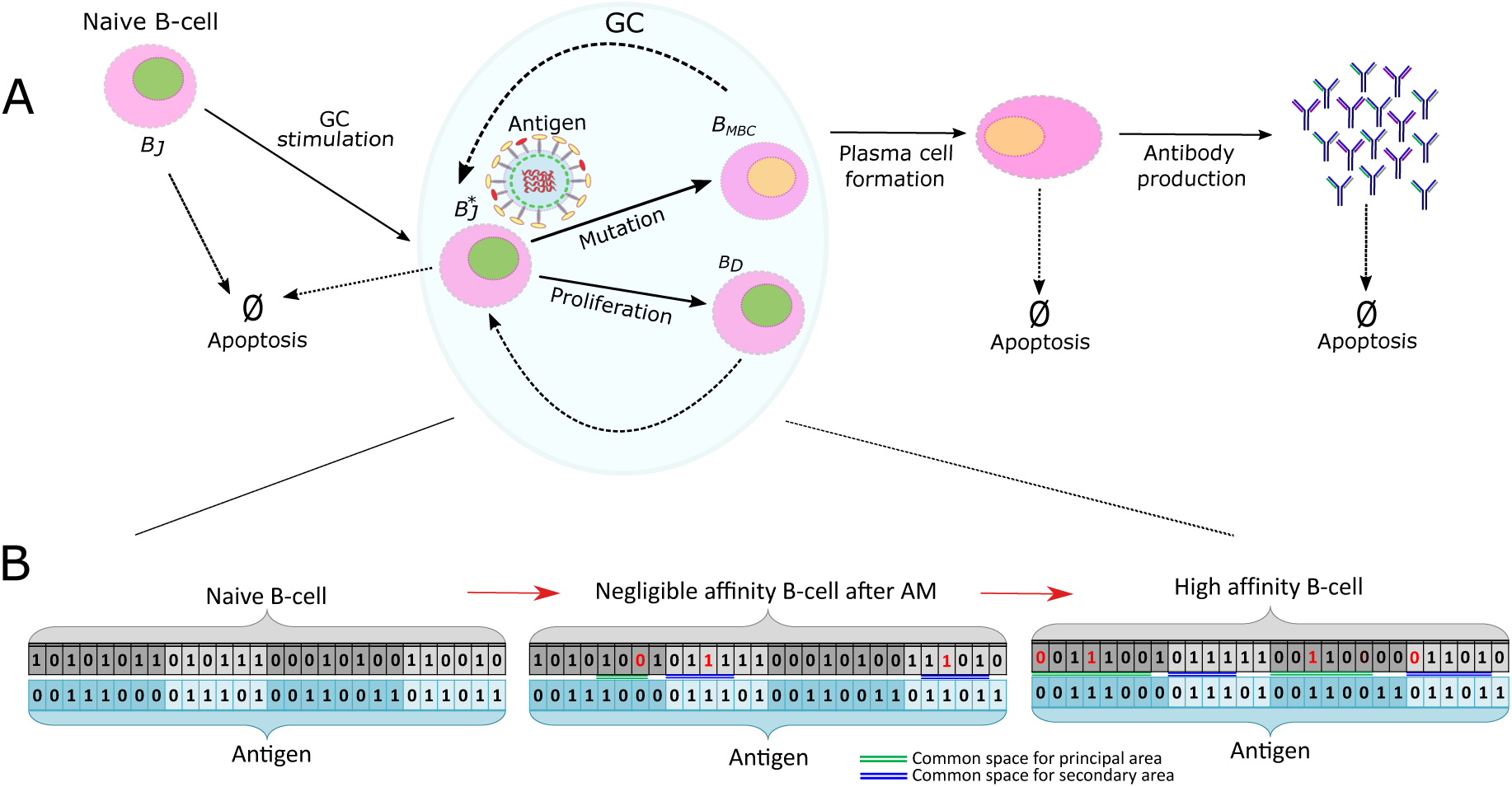
Schematic of the *in-silico* immune system stochastic model. A) The principal immune components of the model following the stimulation of naive B-cells, the AM process to increase fitness, and the production of PCs and Abs. Importantly, each element in the model represent a population complete population and, in the case of PC and Abs, the diversity of clones in the population. Dashed arrows in the GC oval stand for the several cycles of mutation and proliferation to reach affinity for the antigen. B) The principal stages of the AM process, this corresponds to the Fig. 6.

### Antigen and B-cell fitness

For each antigen and B-cell pair, following (15), the matching score indicates how strong the interaction is between them. The binary digits from the antigens and B-cells strings may be analogously thought as the binding sites between them (18). The matching score is determined by three indicators easily identified in Fig 6. 1) the principal matching area, which are the first 8 darker-colored string digits from left to right, 2) the secondary matching area given by the leftover lighter-colored digits and 3) the common space between strings, highlighted in green for common space in the principal area and blue in the secondary matching area. Note that these indicators are separately applied for the x- and y-strings. The matching score (*M*(*B_i_, A_j_*)) for an antigen (*A_j_*) and B-cell (*B_i_*) pair is given as follows

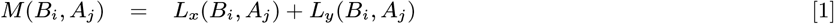

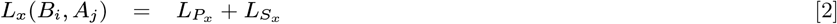

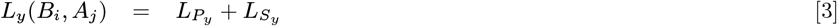

where *L_P_* is the longest common space (consecutive bit-alignments) of the principal area of either x- or y- axis. Similarly, *L_s_*stands for the longest common space in the secondary area of any of the axis, separately. The common space substrings are depicted in Figure6-B,C. *L_P_* and *L_S_* feature the following restriction

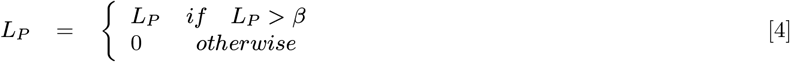

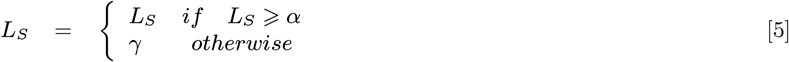

where *α* and *β* are the thresholds above which an antigen and B-cell pair have a favorable match score in the secondary and principal matching areas, respectively. *γ* is a parameter that contributes to the repertoire diversity, avoiding strain-specific behavior, especially during the first CG cycles. The fitness of a B-cell for antigen (*F*(*B_i_*)) is exponentially influenced by the matching score of a B-cell for all antigens and considers the total number of antigens per infection (viral population) (*N_A_*) in the following form

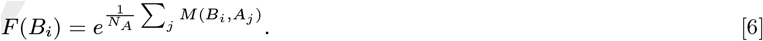

The fitness of a B-cell is higher if its match score against all viruses is high. Here, we assume a relation of B-cells fitness with Abs production, those B-cells that recognize the epitope of an abundant virus in the GC are more likely to receive a survival signal to leave the GC and continue circulating in the system (5, 15).

### B-cell replication, death and affinity mutation

During a GC cycle, B-cells are selected according to its affinity for antigen, the rule of selection is a fitness threshold given by the average fitness of the B-cell population. Those B-cells above the fitness threshold are the high-affinity B-cells (HBC) which are considered to produce two populations of equal size. The first one, namely the daughter cells, conserve the same characteristics of the HBC. The second one, the mutated B-cells (MBC), undergo affinity mutation, as depicted in Fig. 7. We consider this generative process as the replication of B-cells. For each daughter and MBC cells produced, a B-cell is uniformly selected at random to die regardless of its fitness, this allows to keep a constant population of B-cells.

The AM process allows B-cells to modify their string structure to gradually gain affinity for the antigen (15, 17). During AM, a digit of each B-cell strings of the MBC population is uniformly randomly selected to undergo a binary change, this process occurs once per CG cycle. Binary changes, seen as point mutations, make a B-cell to increase the common space of any, the principal or secondary matching area, and therefore to gain fitness. As the binary changes are done at random positions, the risk of reducing the matching score of a B-cell due to AM is latent. Thus, AM is widely responsible for higher or lower matching scores and fitness. In Fig. 6 the AM process is highlighted with red binary digits representing mutations over several CG cycles coming from a naive B-cell to a high affinity one. In the case of Fig. 6-B, there is not an effective matching score due to the short level of bit-alignments for the x-axis and only a few common positions for the secondary y-axis, which is considered as a negligible matching score. In Fig. 6-C, note that after two mutations on the x-axis, the principal matching area has a complete bit-alignments of the string. A similar increment is for the y-axis, forming a HBC.

### Plasma cells and antibodies production

Each HBC has a 50% probability of differentiating into a plasma cell (PC) or a memory B-cell (16), which follows a discrete subset of HBC residing in the GC (21). Memory cells are not considered as acting members of the immune system model since they possess a long lifespan of months to years (16, 22), however, we aim to get insights of the short-term (weeks) activity of PCs and Abs. The PC population decay according to its half-life in a discrete fashion, that is, for every GC cycle, a proportion of the initial PC population is eliminated based on a death probability, forming the PC dynamics shown in Fig. 2-C. This approach considers the tau-leaping methodology (23). Importantly, in a GC cycle, the newly produced PCs are added to the remaining population of previous cycles. Antibodies are produced as copies of PCs and, similarly to the PC population, Abs population raise and decay follow discrete events on every GC cycle. The Abs population decay according to its half-life, depicted Fig 2-D. Eliciting Abs is a direct result of B-cells affinity to antigen, the probabilistic nature of selecting B-cells to become PCs, and the stochasticity involved in both processes. Besides, the modeling mechanisms allow to early appear high-affinity antibodies and a large number of derived plasma cells (24) as depicted in Fig 2-C,D.

### Parameterization

The GC cycle (B-cell division time) is 6 h (5), the B-cell mutation rate is 10^−2^/B-cell/GC cycle (15), the plasma cell half-life is 3 d (13), the antibody half-life is 10 d (16, 25), and the antibody generation is 5 Abs/PC/GC cycle (13, 16). This parameter adjusts the magnitude of Abs response but does not affect the breadth phenomena. On the other hand, the threshold values for the secondary and principal matching areas, *α* and *β*, respectively, serve as tuning parameters to control the breadth outcome, see results for further detail. Finally, the longitude of the binary strings (14 digits) directly comes from the binary-strain-maps in Fig 5.

### High-Performance Computing

Due to computational and efficiency limitations, for instance, one simulation can take up to 48 h to be completed in a modern desktop computer, we employ a High-Performance Computing (HPC) cluster for our simulations, the cluster FUCHS-CSC from the Center for Scientific Computing (Frankfurt, Germany). The cluster is based on 72 dual-socket AMD Magny-Cours CPU compute nodes with 64 GB of RAM, 250 dual-socket AMD Istanbul compute nodes with 32 GB of RAM and 36 quad-socket AMD Magny-Cours compute nodes with 128 GB of RAM each. A simulation is conformed by two parts, one part performing from B-cell generation till PCs formation and the other part is for the Abs production. The first part takes ca. 35 minutes and the second ca. 15 minutes, both in the HPC cluster. The project is coded in Python.

## ACKNOWLEDGMENTS

We thank Raffael Nachbagauer and Florian Krammer (Icahn School of Medicine at Mount Sinai, NY, USA) for providing the data and for valuable comments for the enhancement of this paper. We also thank Franziska Matthäus for her comments. We would like to thank the support of the Alfons und Gertrud Kassel-Stiftung and the Deutsche Forschungsgemeinschaft through the project HE7707/5-1. We also would like to thank the Center for Scientific Computing of the Goethe University Frankfurt for facilitating the FUCHS cluster to operate the simulations.

## References

1. WHO, Influenza (seasonal). World Heal. Organ, https://www.who.int/en/news-room/fact-sheets/detail/influenza-(seasonal) (2018).

2. F Krammer, The human antibody response to influenza a virus infection and vaccination. Nat. Rev. Immunol., 1 (2019).

3. DJ Smith, et al., Mapping the antigenic and genetic evolution of influenza virus. science 305, 371–376 (2004).

4. K Koelle, S Cobey, B Grenfell, M Pascual, Epochal evolution shapes the phylodynamics of interpandemic influenza a (h3n2) in humans. Science 314, 1898–1903 (2006).

5. GD Victora, MC Nussenzweig, Germinal centers. Annu. review immunology 30, 429–457 (2012).

6. NS De Silva, U Klein, Dynamics of b cells in germinal centres. Nat. reviews immunology 15, 137 (2015).

7. M Kuraoka, et al., Complex antigens drive permissive clonal selection in germinal centers. Immunity 44, 542–552 (2016).

8. A Amitai, L Mesin, GD Victora, M Kardar, AK Chakraborty, A population dynamics model for clonal diversity in a germinal center. Front. microbiology 8, 1693 (2017).

9. J Lee, et al., Persistent antibody clonotypes dominate the serum response to influenza over multiple years and repeated vaccinations. Cell host & microbe 25, 367–376 (2019).

10. F Krammer, P Palese, Influenza virus hemagglutinin stalk-based antibodies and vaccines. Curr. opinion virology 3, 521–530 (2013).

11. R Nachbagauer, et al., Defining the antibody cross-reactome directed against the influenza virus surface glycoproteins. Nat. immunology 18, 464 (2017).

12. KR McCarthy, et al., Memory b cells that cross-react with group 1 and group 2 influenza a viruses are abundant in adult human repertoires. Immunity 48, 174–184 (2018).

13. DJ Smith, S Forrest, DH Ackley, AS Perelson, Variable efficacy of repeated annual influenza vaccination. Proc. Natl. Acad. Sci. 96, 14001–14006 (1999).

14. PA Robert, AL Marschall, M Meyer-Hermann, Induction of broadly neutralizing antibodies in germinal centre simulations. Curr. opinion biotechnology 51, 137–145 (2018).

15. S Luo, AS Perelson, Competitive exclusion by autologous antibodies can prevent broad hiv-1 antibodies from arising. Proc. Natl. Acad. Sci. 112, 11654–11659 (2015).

16. S Chaudhury, J Reifman, A Wallqvist, Simulation of b cell affinity maturation explains enhanced antibody cross-reactivity induced by the polyvalent malaria vaccine ama1. The J. Immunol. 193, 2073–2086 (2014).

17. S Wang, et al., Manipulating the selection forces during affinity maturation to generate cross-reactive hiv antibodies. Cell 160, 785–797 (2015).

18. S Khailaie, PA Robert, A Toker, J Huehn, M Meyer-Hermann, A signal integration model of thymic selection and natural regulatory t cell commitment. The J. Immunol., 1400889 (2014).

19. A Lapedes, R Farber, etal., The geometry of shape space: application to influenza. J. theoretical biology 212, 57–70 (2001).

20. K Murphy, C Weaver, Janeway’s immunobiology. (Garland Science), (2016).

21. NJ Kräutler, et al., Differentiation of germinal center b cells into plasma cells is initiated by high-affinity antigen and completed by tfh cells. J. Exp. Medicine 214, 1259–1267 (2017).

22. I Dogan, et al., Multiple layers of b cell memory with different effector functions. Nat. immunology 10, 1292 (2009).

23. Y Cao, DT Gillespie, LR Petzold, Efficient step size selection for the tau-leaping simulation method. The J. chemical physics 124, 044109 (2006).

24. M Meyer-Hermann, et al., A theory of germinal center b cell selection, division, and exit. Cell reports 2, 162–174 (2012).

25. P Vieira, K Rajewsky, The half-lives of serum immunoglobulins in adult mice. Eur. journal immunology 18, 313–316 (1988).

26. JM Tas, et al., Visualizing antibody affinity maturation in germinal centers. Science 351, 1048–1054 (2016).

27. AG Schmidt, et al., Viral receptor-binding site antibodies with diverse germline origins. Cell 161, 1026–1034 (2015).

28. D Angeletti, et al., Defining b cell immunodominance to viruses. Nat. immunology 18, 456 (2017).

29. GD Victora, PC Wilson, Germinal center selection and the antibody response to influenza. Cell 163, 545–548 (2015).

30. D Lingwood, et al., Structural and genetic basis for development of broadly neutralizing influenza antibodies. Nature 489, 566 (2012).

31. SF Andrews, et al., Immune history profoundly affects broadly protective b cell responses to influenza. Sci. translational medicine 7, 316ra192–316ra192 (2015).

32. CS Anderson, et al., Natural and directed antigenic drift of the h1 influenza virus hemagglutinin stalk domain. Sci. reports 7, 14614 (2017).

33. E Kirkpatrick, X Qiu, PC Wilson, J Bahl, F Krammer, The influenza virus hemagglutinin head evolves faster than the stalk domain. Sci. reports 8, 10432 (2018).

34. MD Pauly, MC Procario, AS Lauring, A novel twelve class fluctuation test reveals higher than expected mutation rates for influenza a viruses. Elife 6, e26437 (2017).

35. KE Johnson, T Song, B Greenbaum, E Ghedin, Getting the flu: 5 key facts about influenza virus evolution. PLoS pathogens 13, e1006450 (2017).

36. EC Holmes, et al., Whole-genome analysis of human influenza a virus reveals multiple persistent lineages and reassortment among recent h3n2 viruses. PLoS biology 3, e300 (2005).

37. BL Tesini, et al., Broad hemagglutinin-specific memory b cell expansion by seasonal influenza virus infection reflects early-life imprinting and adaptation to the infecting virus. J. virology, JVI–00169 (2019).

38. K Ito, et al., Gnarled-trunk evolutionary model of influenza a virus hemagglutinin. PLoS One 6, e25953 (2011).

39. JM Fonville, et al., Antibody landscapes after influenza virus infection or vaccination. Science 346, 996–1000 (2014).

40. ST Liu, et al., Antigenic sites in influenza h1 hemagglutinin display species-specific immunodominance. The J. clinical investigation 128 (2018).

