## Supplemental for "Uncovering antibody cross-reaction dynamics in influenza A infections"

### 1. Two's complement and bit-space

To convert a binary number to two's complement we revert each bit of this number, meaning 1 changes to 0 and 0 changes to 1, and then add 1 to the result. For positive numbers the leftmost bit is 0 and for negative numbers the leftmost bit is 1. The number zero has only one representation (in contrary to one's complement).

In the binary-strain map, as the Cartesian fashion, it covers the x-axis and y-axis where the axis range is given by $(-\frac{2^n}{2}, +\frac{2^n}{2} - 1)$, where $n \in \mathbb{N}$ is the range-tuning parameter (bit resolution). The reason to subtract the value of 1 from the positive side is to take into account the zero in the bit map, see Fig. (S1). The parameter $n$ must be adjusted to set a covering space that is sufficiently big to capture the coordinates range of the antigenic/genetic map. If $n = 10$, for instance, the x- and y-axis would cover the range $(-512, +511)$. The binary representation of each decimal number is the two's complement using a signed integer of size $n$. The positive values representation of a base ten number into binary is direct, just considering the size of $n$. In the case of representing a number with negative value, "$-3$" for instance, considering $n = 4$, it follows three straightforward steps. First give the binary value in positive manner $'0011'$. Next, set the complement of the binary number "1100". This is made applying the *not* logic function. If the character is '1', it is changed to '0' and vice-versa. Finally, add "1" to the complement "1101". Thus, the two's complement binary representation of size 4 of the signed integer $-3$ is 1101.

For each strain cluster centroid in Fig. 1 in the main text, using $n = 14$, we set the centroid binary coordinates, presenting the genetic map coordinates and the corresponding binary values in the binary-strain map. Note that in the binary representation all positive values have the first bit (from left to right) set to "0", conversely, the binary negative values have a "1" in its first position, this bit is the most-significant bit (MSB).

**Sign restriction.** A binary-string with a positive or negative value is distinguished by the most-left bit in the string, the sign-bit. Digits 0 and 1 stand for positive and negative values, respectively. The possibility of finding two almost equivalent strings whose only difference is the sign-bit is high, therefore, there may be strings considered of high affinity because all bits are equivalent to antigen bits, but the sign-bit differs. This issue affects B-cells proliferation since the high affinity of a string with only the sign-bit different may be selected to reproduce. We avoid this issue by setting the initial B-cell repertoire with 0 in the sign-bit and restricting this bit to mutation, meaning that it can never take the value of 1. The principal test follows these conditions.

**Parameter $\gamma$.** The parameter $\gamma = 2$ is for avoiding strain-specific behavior. This parameter contributes to the repertoire diversity since those B-cells with a negligible affinity still may be able to compete for antigen, especially during the first GC cycles.

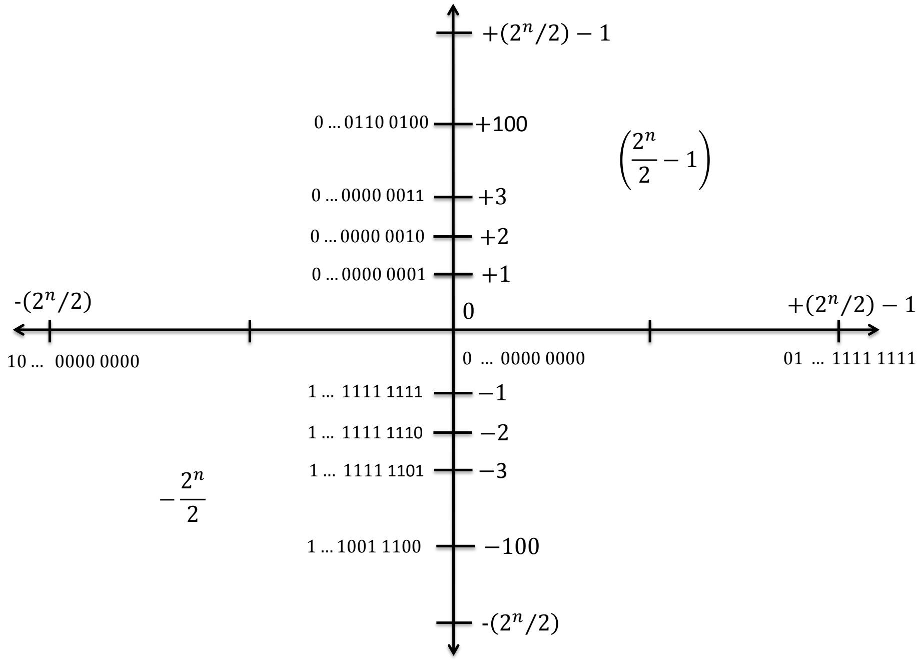


**Fig. (S1)** Bit-space using two's complement binary numbers. For any value of $n \in \mathbb{N}$, the positive side of x- and y-axis of the bit-space goes from zero to $\frac{2^n}{2} - 1$ while the negative side of x- and y-axis goes from -1 to $-\frac{2^n}{2}$. Depending of the value of $n$, the resolution of bit-space can cover more 10-base numbers whose values are represented by binary strings. Note that the first digit of the strings in the negative axes is "1". on the other hand, the first digit of the strings in the positive axes is "0".

### 2. H1N1 response and cross-reaction

Following the Nachbagauer experiments for cross-reactive Abs induction, For H1N1 infections, the first infection is made with a human seasonal strain A/New Caledonia/20/1999 (NC99), followed by the 2009 human pandemic H1N1 strain A/Netherlands/602/2009 (NL09) six weeks later. NL09 is an isolate antigenically identical to the prototype pandemic H1N1 strain A/California/04/09 (Cal09). The *in-silico* experiment in Fig. 1-D-F (main text) follows the mice experiments and employs 10 simulations of the profile described above.

Fig. (S2). shows the H1N1 strains space and dynamics of B-cells, plasma cells and antibodies of one of ten simulations. Fig. (S2-A) is a snapshot of the resulting distribution of B-cells 42 days post-infection (dpi) after the second infection. Plasma cells and Abs dynamics differences are closely related to each population's half-life.

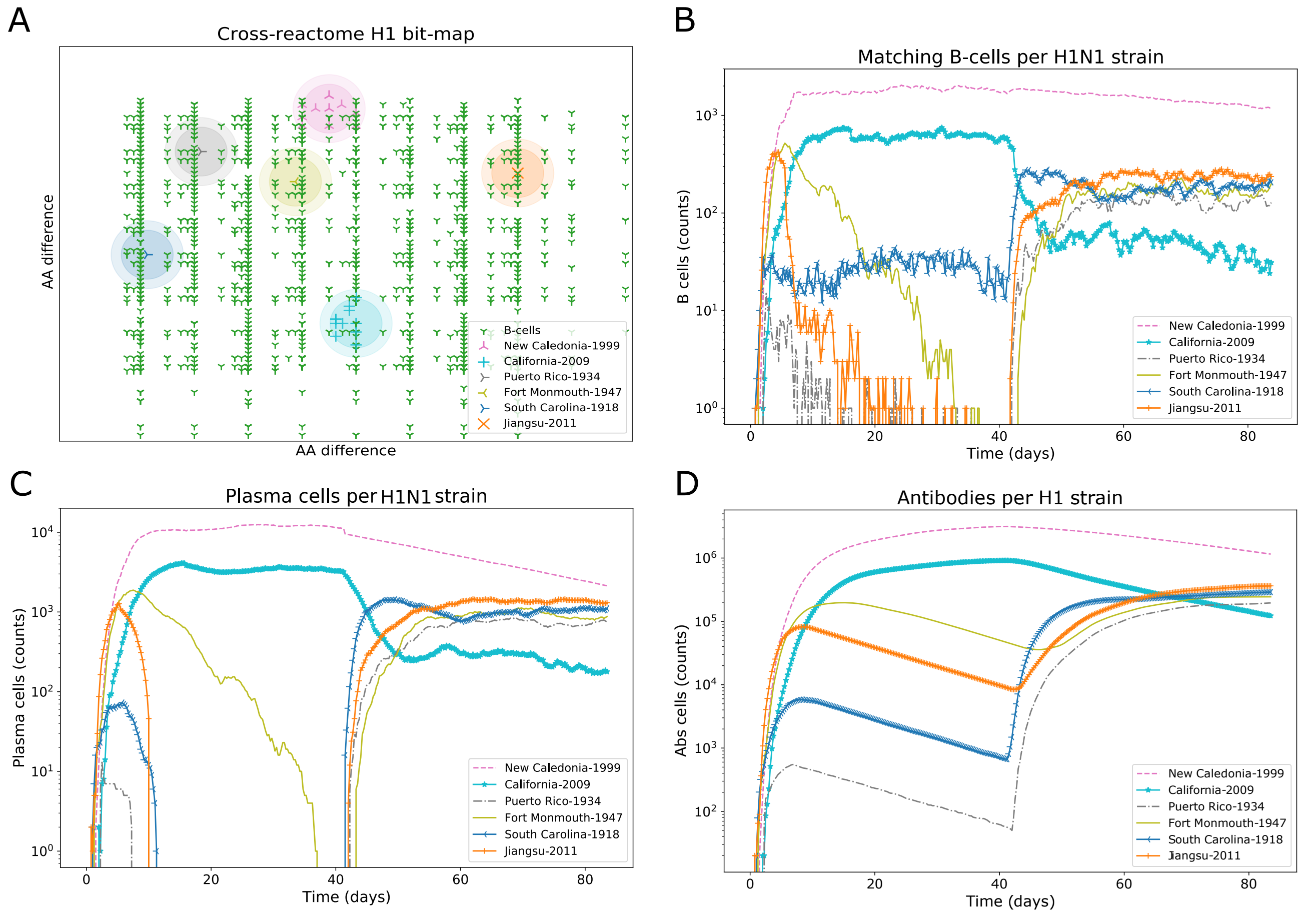


**Fig. (S2)** Influenza H1N1 strains *shape-space* and immune system components dynamics. (A) The genetic binary-map of Fig. 1-B in the man text is used to show the B-cell population covering most of the H1N1 influenza strains clusters after the Cal09 infection. The initial population of naive B-cells covers the entire genetic binary-map, however, as a result of AM, the B-cell population targets the H1N1 strains space. (B) A B-cell that falls into the diameter of a strain cluster is considered as a matching B-cell for that strain. The quantity of B-cells inside each cluster is plotted respect to time. (C) A high-affinity B-cell has the chance to generate a plasma cell which decay according to its half-life. Due to a combined effect of the high threshold for the principal area during the second infection and AM, the plasma cells in the Cal09 infection manage to remain more time than for the NC99 infection. (D) The antibody population for each strain is generated by plasma cells targeting that corresponding strain. The four panels (A-D) are the outcome of one of the 10 performed simulations in the principal test.

### 3. Affinity threshold in secondary area, $\alpha = 3$

Fig. (S3) depicts the outcome of a fixed threshold for the secondary area, $\alpha = 3$, in the binary strings. Having this threshold has implications in the cross-reaction magnitude, especially during the second infection. Note that the resulting phylogenetic trees and Abs dynamics in the second infection, for both subtypes, inherits a strain-specific behavior from the first infection, which does not correspond with the experimental data.

For both influenza strains in Fig. (S3), using a fixed $\alpha = 3$ for the first and second infections, an strain specific framework results especially during the second infection. The phylogenetic trees of the first infection in Fig. (S3-A,D) for both influenza strains are naturally equivalent to those in Fig. 1-A,D. On the other hand, there are clear differences between the phylogenetic trees of the second infection in Fig. (S3-B), E and those in Fig. 1-B,E. An outstanding difference refers to the high level of Abs that map to darker heat-map colors in both, H3N2 and H1N1 strains. The Vic11 and Cal09 develop high Abs levels compared to the low reactivity of the other strains which feature clearer colors in the heat-map. Note that in an H3N2 simulation, only Phil82, Vic11 and Mass11 strains develop high counts of Abs, shown in Fig. (S3-B,C), compared to the all-strains Abs response in Fig. 1-C. Also, an H1N1 simulation in Fig. (S3-F) shows a particular outcome of Cal09 Abs while in Fig. 1-F, all of the H1N1 strains develop high levels of Abs.

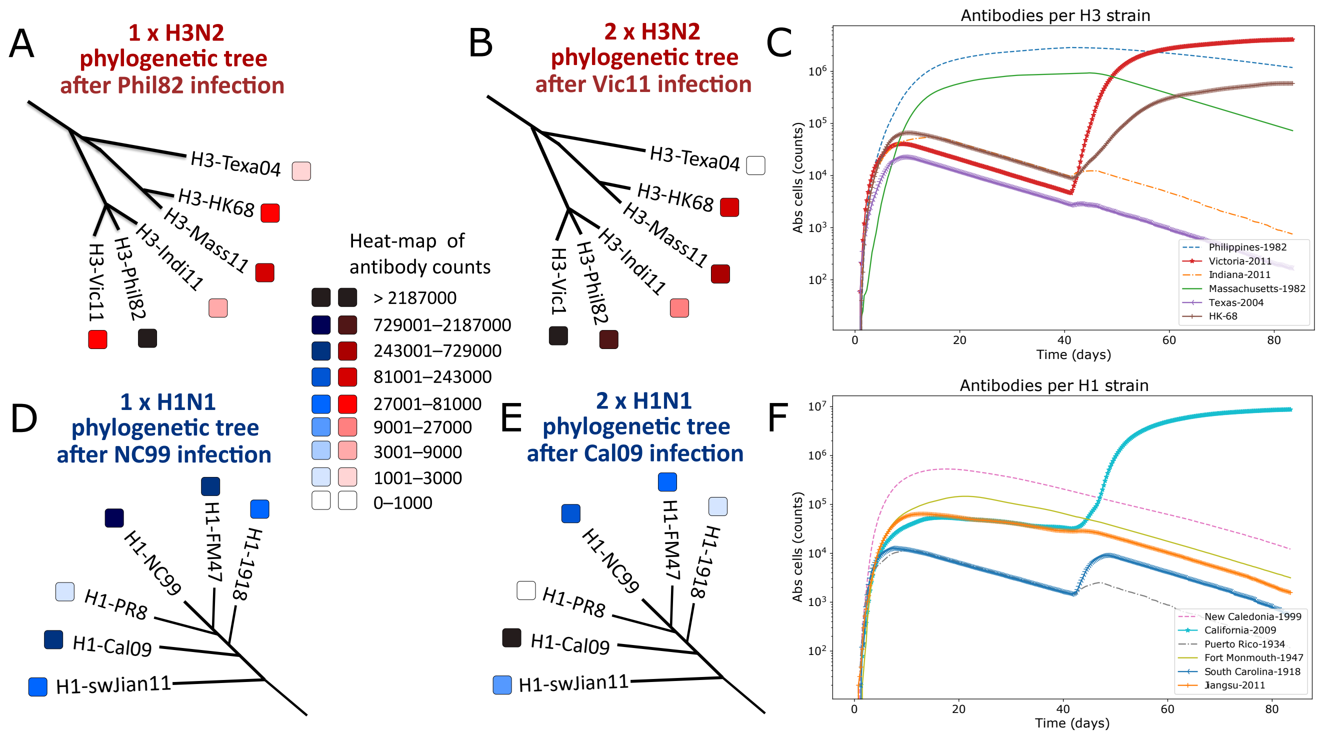


**Fig. (S3)** Test of a 3-bit affinity threshold for secondary area. (A-B) A red-colored heat-map display the intervals of Abs counts for different H3N2 strains with the same characteristics of Fig. 3 of the main text. (A) Abs counts after the first influenza infection with Philippines-1982 (Phil-82) strain. (B) Abs counts after the second infection with Victoria-2011 (Vic-11) strain. (C) The dynamics of Abs for both H3N2 infections in time with each H3 strain is depicted with different colors and symbols. (D-E) A blue-colored heat-map stand for the intervals of Abs counts for one or two consecutive H1N1 infections with the same characteristics of Fig. 3 of the main text. (D) The first H1N1 infection is with the New Caledonia-1999 strain (NC99) followed by H1N1 isolate Netherlands-09 (NL09) (E). (F) The dynamics of Abs counts for both H1N1 infections in time (days), each H1N1 strain is depicted with different colors and symbols. The Abs counts represent the result of 10 simulations for H3N2 or H1N1, separately. The panorama between infections with fixed affinity threshold, $\alpha = 3$, shows a remarked strain-specific activity during both infections. This effect leads to the marginal cross-reaction outcome during the second infection. Note that the first infection in both subtypes is consistent with the principal experiment in Fig. 3-A,D.

### 4. Affinity threshold in secondary area, $\alpha = 4$

Fig. (S4) depicts the outcome of a fixed threshold for the secondary area, $\alpha = 4$, in the binary strings. During the first infection, although the breadth of the Abs cross-response is wide, a weak specificity for the strain of infection is present. This influence is inherited after the second infection. Note that the magnitude of Abs response in the phylogenetic trees stays in the medium level of the heat-map in both infections of both subtypes. The test of a fixed $\alpha = 4$ for the first and second infections of both strains, H3N2 and H1N1, promote a low-affinity framework during the first infection, however, this behavior is inherited to the second infection. The results of this test are shown in Fig. (S4).

The phylogenetic trees of the first infection in Fig. (S4-A,D) for both influenza strains report that lower levels of Abs are developed for the strains of the infection, Phil82 and NC99, respect to other strains such as the Indi11 or FM47, H3N2 and H1N1, respectively. This phenomenon is also observed in the Abs dynamics in Fig. (S4-C,F) where during the first infection time window, the Phil82 and NC99 strains remain below most of the other H3 or H1 strains. During the second infection, a similar result can be observed.

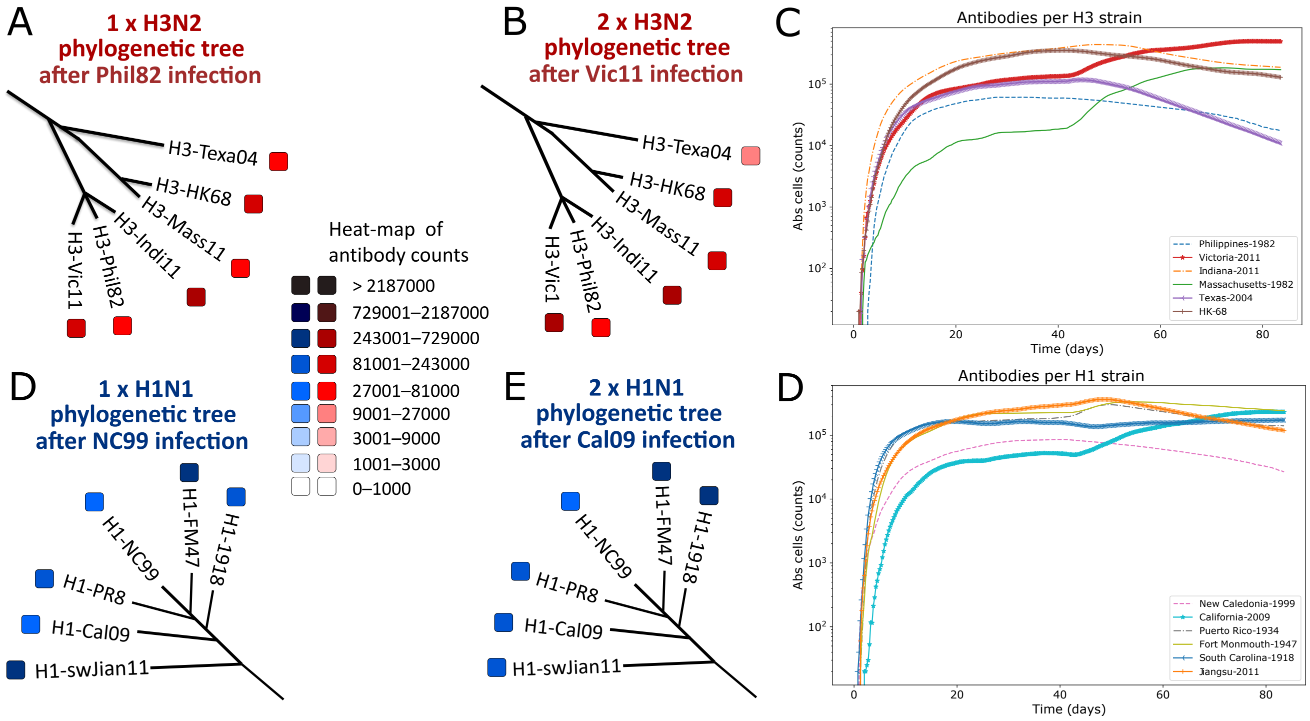


**Fig. (S4)** Test of a 4-bit affinity threshold for secondary area. (A-B) A red-colored heat-map display the intervals of Abs counts for different H3N2 strains with the same characteristics of Fig. 3 of the main text. (A) Abs counts after the first influenza infection with Philippines-1982 (Phil-82) strain. (B) Abs counts after the second infection with Victoria-2011 (Vic-11) strain. (C) The dynamics of Abs for both H3N2 infections in time with each H3 strain is depicted with different colors and symbols. (D-E) A blue-colored heat-map stand for the intervals of Abs counts for one or two consecutive H1N1 infections with the same characteristics of Fig. 3 of the main text. (D) The first H1N1 infection is with the New Caledonia-1999 strain (NC99) followed by H1N1 isolate Netherlands-09 (NL09) (E). (F) The dynamics of Abs counts for both H1N1 infections in time (days), each H1N1 strain is depicted with different colors and symbols. The Abs counts represent the result of 10 simulations for H3N2 or H1N1, separately. The panorama between infections with fixed affinity threshold, $\alpha = 4$, shows a remarked cross-reactivity scheme during both infections. This effect leads to low strain affinity for the first infection, a scheme that is inherited for the second one. Although the breadth of Abs response is positive, the magnitude of Abs outcome is considerably lower compared to the principal test in Fig. 3 of the main text. This may have further implications in Abs and memory B-cells limiting protection of future infections which is not fully consistent with experiments.

### 5. Internal and external clusters testing

The influence of clusters genetic similarity is tested through consecutive infections of closely related antigen clusters (internal clusters) and genetically distanced clusters (external clusters). Fig. (S5) and Fig. (S6) show the results for internal and external clusters, respectively. The H3N2 internal clusters are Mass11 and HK68 for first and second infection, respectively. For H1N1, the internal clusters are NC99 and FM47 for first and second infection, respectively.

Results with the internal clusters in Fig. (S5-A,B) report a similar outcome to the principal test, high Abs levels are developed, especially during the second infection. The Fig. (S5-G-I) portraits the H3N2 binary-maps with the B-cells covering the corresponding area of each strain cluster for the Mass11 and HK68 strains, respectively. For the H1N1 strains, the first infection behaves with a strain-specific tendency, similar to the principal test of the H1N1 strain. The breadth and magnitude of the second H1N1 infection develop Abs to all of the H1N1 strains in Fig. (S5-E-F). The distribution of B-cells for both H1N1 infections is shown in Fig. (S5-J-K). Of note, the B-cells shape of the genetic binary-map in Fig. (S5-J) is equivalent of the principal H1N1 test in Fig 1-D, F, since the first infection in both tests is with the NC99 strain.

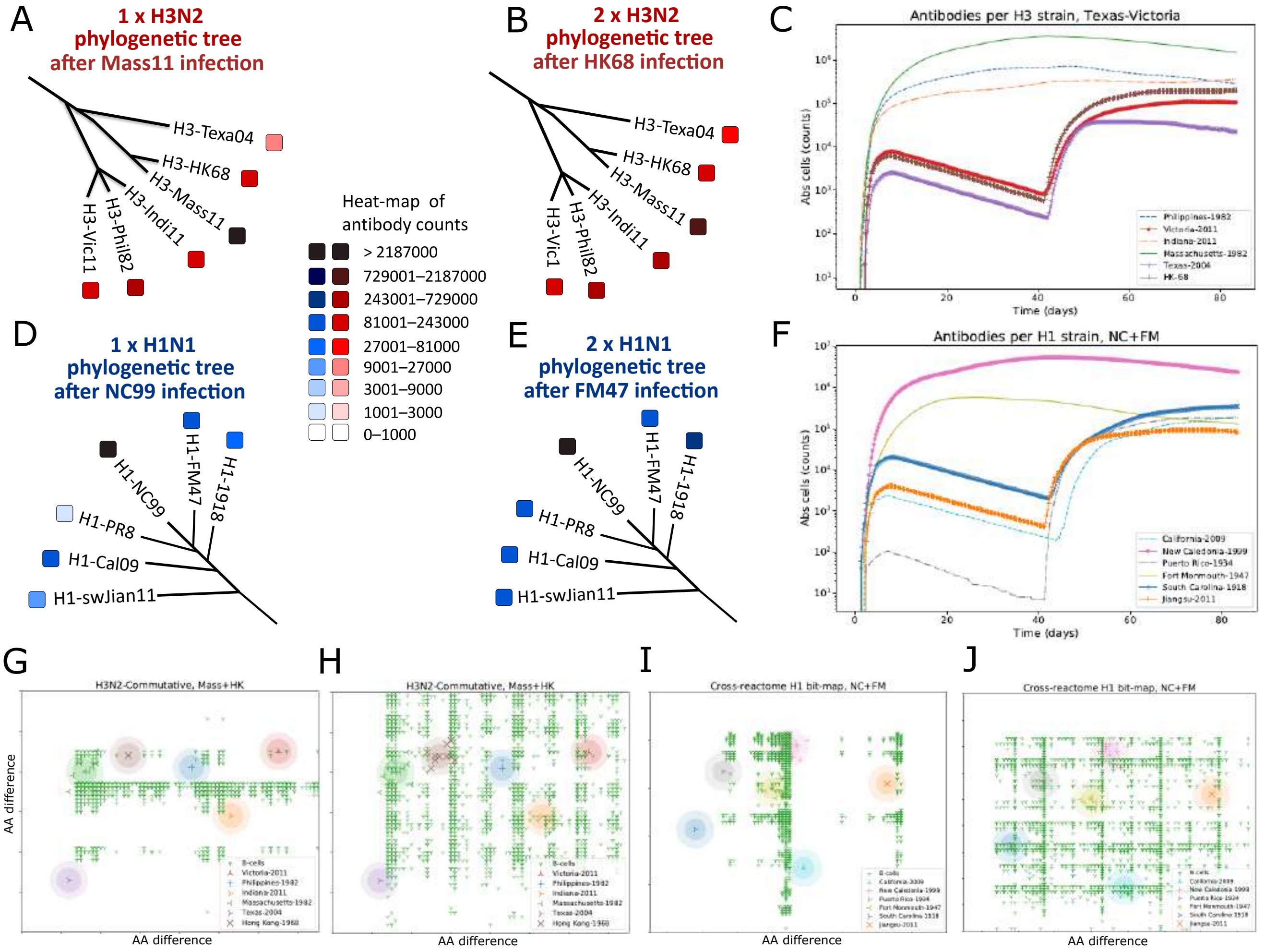


**Fig. (S5)** The internal clusters test shows the influence of genetic similarity between clusters through consecutive infections of closely related antigen clusters (internal clusters). (A-B) A red-colored heat-map display the intervals of Abs counts for different H3N2 strains with the same characteristics of Fig. 3 of the main text. (A) Abs counts after the first influenza infection with Mass11. (B) Abs counts after the second infection with HK68 strain. (C) The dynamics of Abs for both H3N2 infections in time with each H3 strain is depicted with different colors and symbols. (D-E) A blue-colored heat-map stand for the intervals of Abs counts for one or two consecutive H1N1 infections with the same characteristics of Fig. 3 of the main text. (D) The first H1N1 infection is with the NC99 followed by FM47 strain (E). (F) The dynamics of Abs counts for both H1N1 infections in time (days), each H1N1 strain is depicted with different colors and symbols. The Abs counts represent the result of 10 simulations for H3N2 or H1N1, separately. B-cell repertory in H3 first and second infections, G-I, respectively. B-cell repertory of H1 first and second infections, J-K, respectively. Slightly different antigens may derive effective cross-reaction in both infections.

Fig. (S6) show the results for external clusters. The H3N2 external clusters are Tex04 and Vic11 for first and second infection, respectively. For H1N1, the external clusters are SC18 and Jiang11 for first and second infection, respectively. An important effect of the strain position is observed in the first H3N2 infection of the external clusters test. The phylogenetic tree, the Abs dynamics, and the B-cells distribution in Fig. (S6-A,C,G), respectively, show a limited cross-reaction to the infection with the Tex04 strain. This result is due to the majority of B-cells are focused on the infection strain. However, the second H3N2 infection with the most distanced Vic11 strain, respect to Tex04, can produce a cross-reaction effect similar to the principal test. Importantly, the H1N1 infection test of the external clusters surprisingly behaves as the principal test. The Abs response portrait of the phylogenetic tree, the Abs dynamics, and the B-cell distribution in Fig. (S6-E,F,J) is equivalent to the test in Fig. 1-E,F (main text) and Fig.(S2-A), respectively.

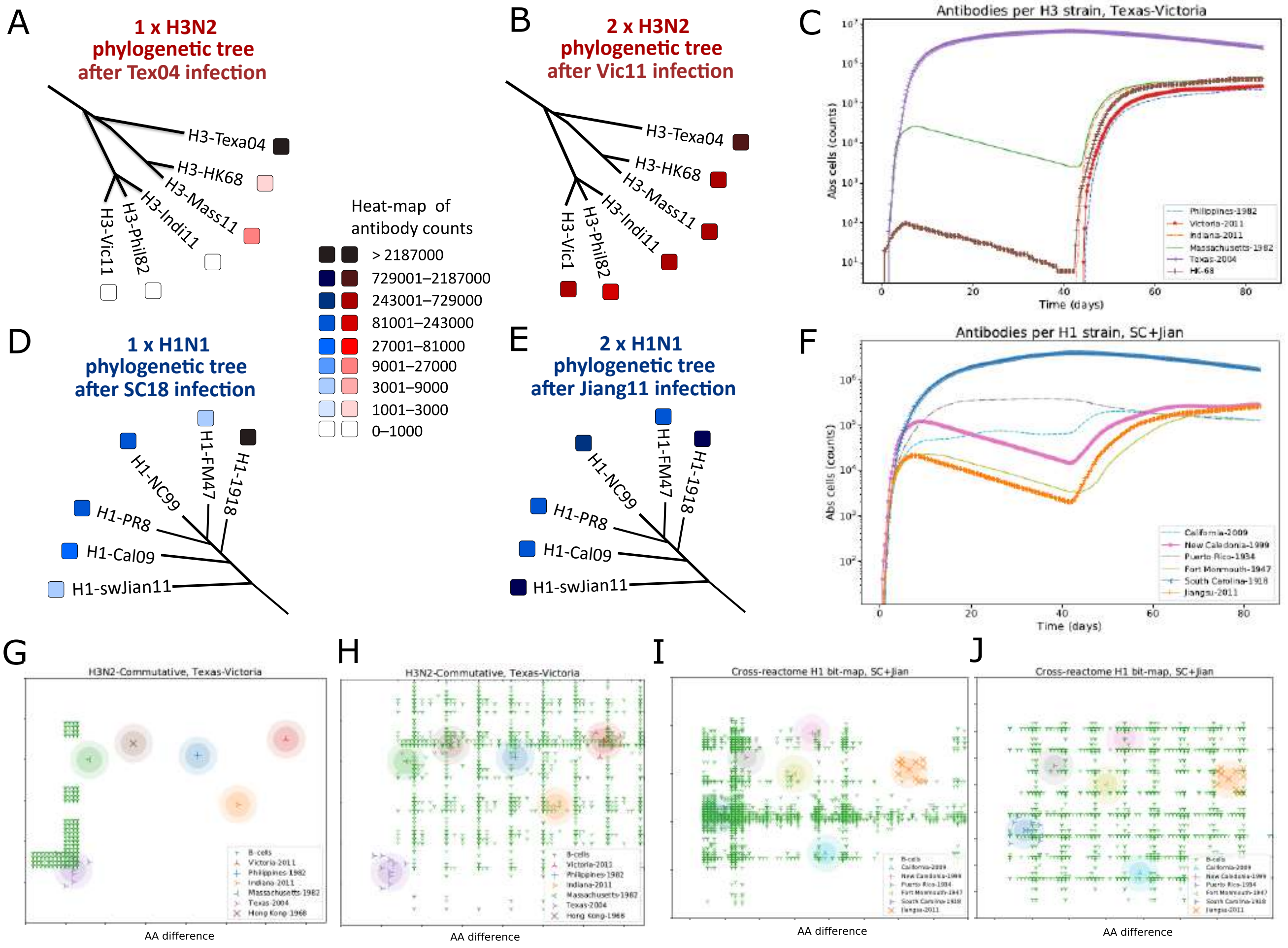


**Fig. (S6)** The external clusters test shows the influence of genetic similarity between clusters through consecutive infections of genetically distanced clusters (external clusters). (A-B) A red-colored heat-map display the intervals of Abs counts for different H3N2 strains with the same characteristics of Fig. 3 of the main text. (A) Abs counts after the first influenza infection with Texo4. (B) Abs counts after the second infection with Vic11 strain. (C) The dynamics of Abs for both H3N2 infections in time with each H3 strain is depicted with different colors and symbols. (D-E) A blue-colored heat-map stand for the intervals of Abs counts for one or two consecutive H1N1 infections with the same characteristics of Fig. 3 of the main text. (D) The first H1N1 infection is with the SC18 followed by Jiang11 strain (E). (F) The dynamics of Abs counts for both H1N1 infections in time (days), each H1N1 strain is depicted with different colors and symbols. The Abs counts represent the result of 10 simulations for H3N2 or H1N1, separately. B-cell repertory in H3 first and second infections, G-I, respectively. B-cell repertory of H1 first and second infections, J-K, respectively. Distanced antigens may (H1) or may not (H3) derive effective cross-reaction in both infections.

### 6. Initial shape and size of naive B-cells

We test the influence of the initial naive B-cells repertory shape and size. We show that newly-generated repertory (principal test) versus a fixed repertory employed by several simulations make weak outcome consequence for breadth and magnitude of Abs. This result is depicted in Fig. S7.

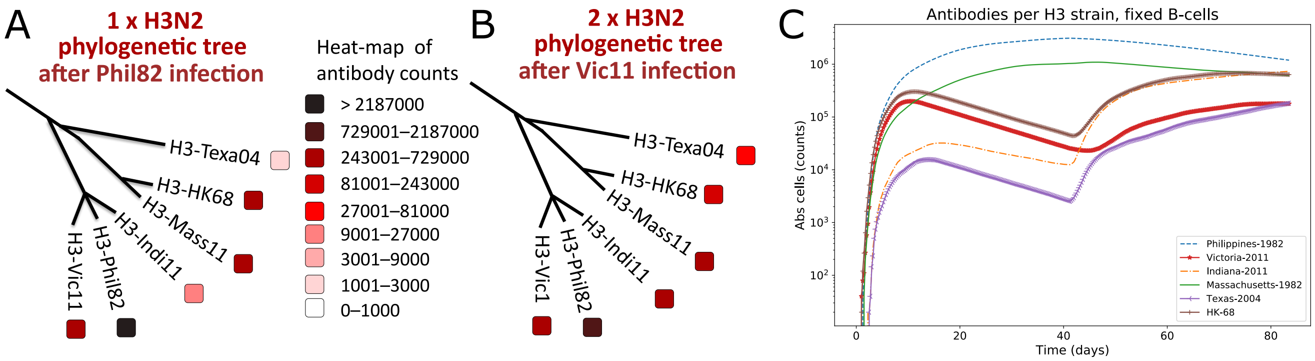


**Fig. (S7)** Fixed naive B-cells test. Newly-generated naive B-cells repertory per simulation (principal test) versus a fixed B-cell repertory employed by several simulations may add to the scheme of a weak consequence of the initial naive B-cells repertory. (A-B) A red-colored heat-map display the intervals of Abs counts for different H3N2 strains with the same characteristics of Fig. 3 of the main text. (A) Abs counts after the first influenza infection with Phil82. (B) Abs counts after the second Vic11 infection. (C) The dynamics of Abs for both H3N2 infections in time with each H3 strain is depicted with different colors and symbols. (D-E) A blue-colored heat-map stand for the intervals of Abs counts for one or two consecutive H1N1 infections with the same characteristics of Fig. 3 of the main text. (D) The first H1N1 infection is with the NC99 followed by Cal09 scheme of infection (E). (F) The dynamics of Abs counts for both H1N1 infections in time (days), each H1N1 strain is depicted with different colors and symbols. The Abs counts represent the result of 10 simulations for H3N2 or H1N1, separately.

We test the influence of the initial naive B-cells repertory shape and size. We show that a repertory from 1000 up to 10 thousand B-cells, proportionally speaking, produce equivalent outcomes. This result is depicted in Fig. S13.

Additionally, we show that different statistic and spatial distributions of the initial B-cell repertory may produce equivalent breadth and magnitude Abs outcomes. This result is depicted in Fig. S14.

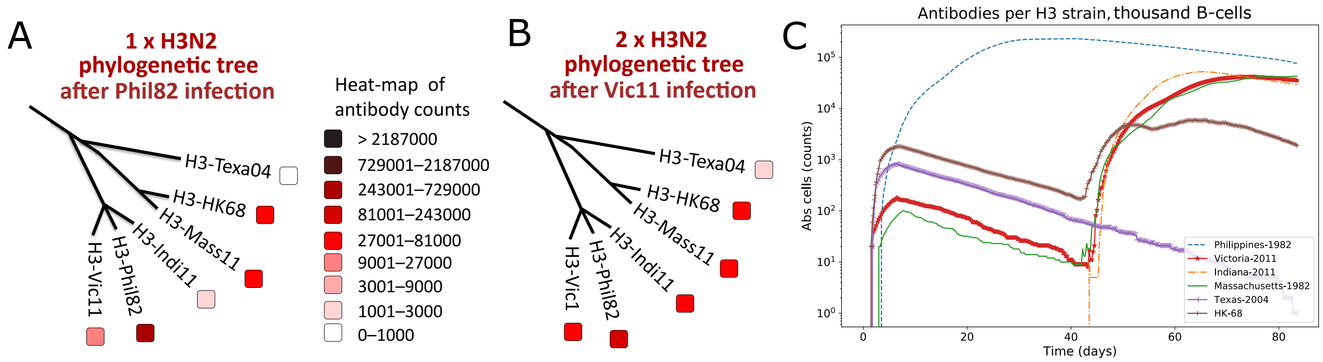


**Fig. (S8)** Test with 1000 naive B-cells. Proportionally speaking, the qualitative scheme of infections may be represented with a repertory of minimum of 1000 naive B-cells.

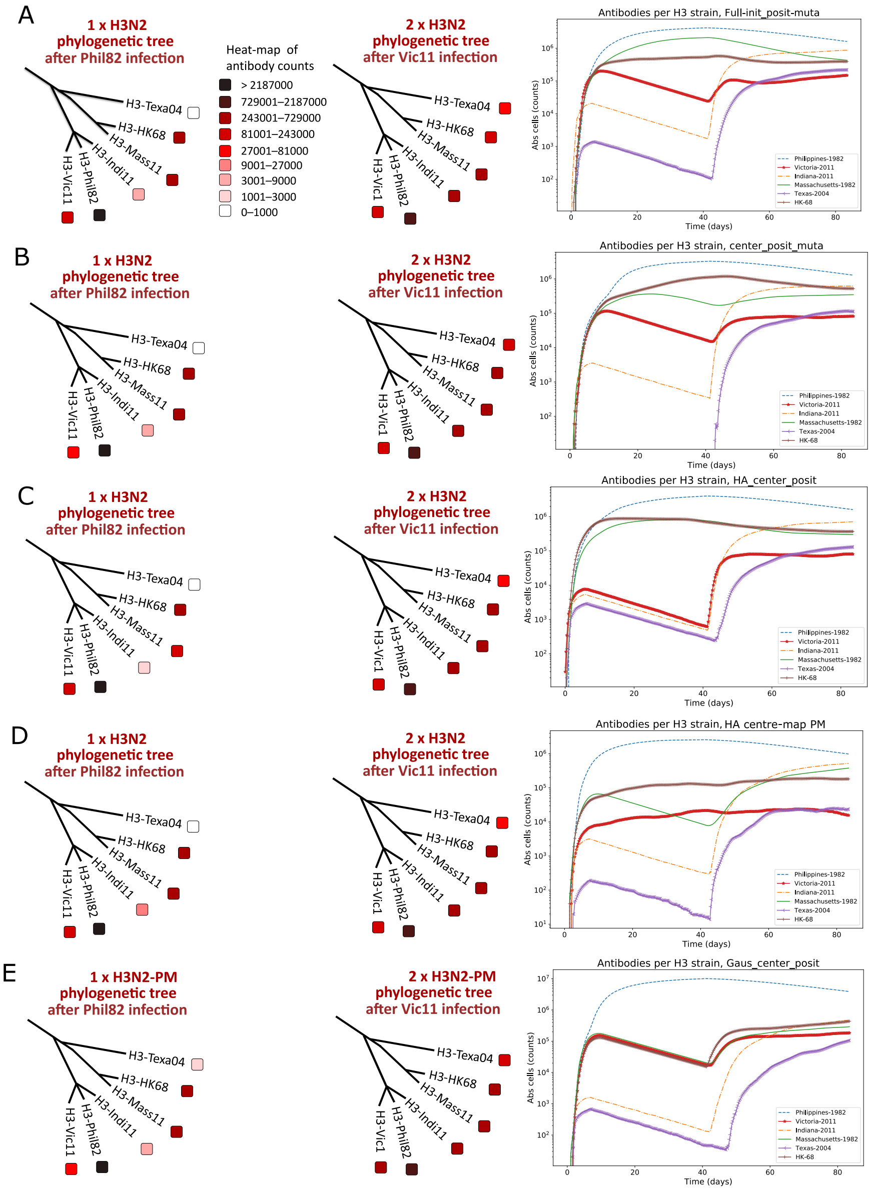


**Fig. (S9)** Test with different statistic and spatial distributions of the initial naive B-cell repertories. Throughout diverse simulation frameworks, the random distribution (uniform and Gaussian), quantity (Fig. S13), and space coverage of initial naive B-cells weakly influence the complete qualitative picture of cross-reactive antibodies. A red-colored heat-map display the intervals of Abs counts for different H3N2 strains with the same characteristics of Fig. 3 of the main text. (A) Consecutive infection with an initial B-cell repertory covering the complete bit-space range of Fig. S1 with uniform distribution. (B) Results of a repertory uniformly distributed but covering only the center of the bit-space. (C) The space coverage is limited to the area where the H3N2 strains are located on the map, with uniform distribution. (D-E) Results of a Gaussian distribution in the center of H3 HAs and the center of the bit-space, respectively

106

### 7. Time windows between infections

Time window of 10 days between infections. A window up to 6 weeks between infections (as in the principal test) manages to develop the best Abs magnitude for the first infection and therefore, a higher response magnitude for the second infection too.

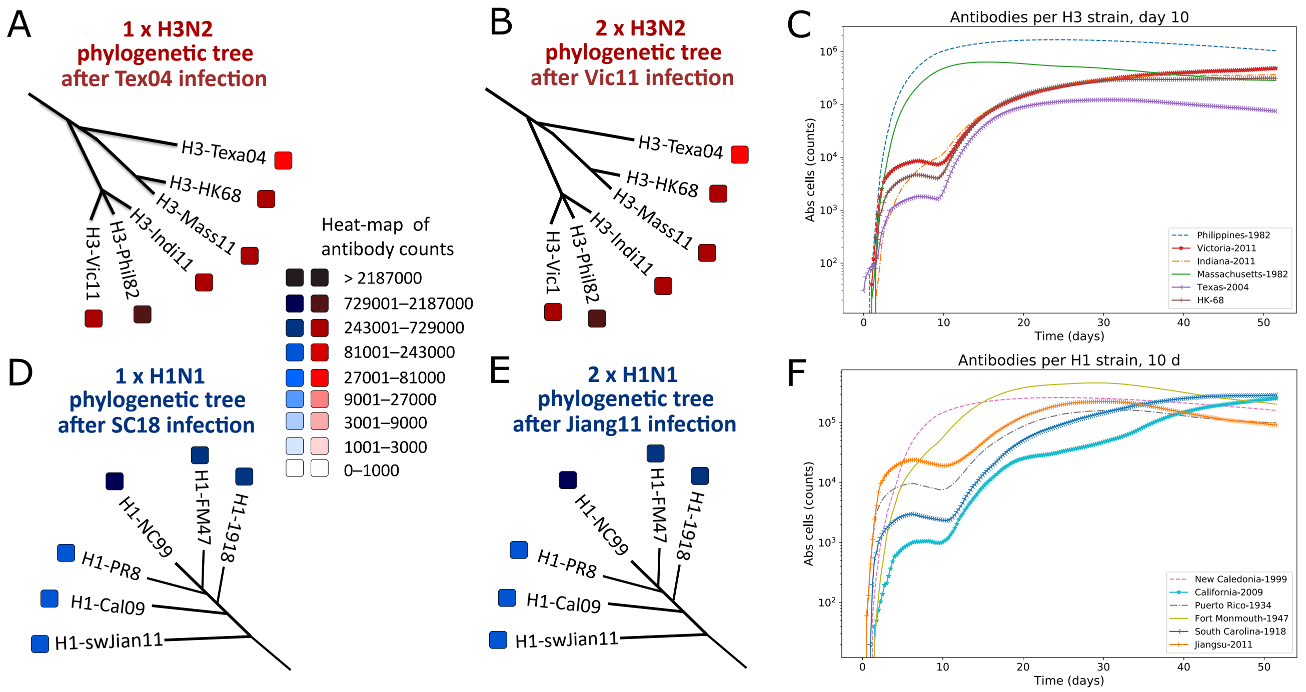


**Fig. (S10)** Test of second infection at 10 days after first infection. (A-B) A red-colored heat-map display the intervals of Abs counts for different H3N2 strains with the same characteristics of Fig. 3 of the main text. (A) Abs counts after the first influenza infection with Phil82. (B) Abs counts after the second infection with Vic11 strain. (C) The dynamics of Abs for both H3N2 infections in time with each H3 strain is depicted with different colors and symbols. (D-E) A blue-colored heat-map stand for the intervals of Abs counts for one or two consecutive H1N1 infections with the same characteristics of Fig. 3 of the main text. (D) The first H1N1 infection is with the NC99 followed by NL09 strain (E). (F) The dynamics of Abs counts for both H1N1 infections in time (days), each H1N1 strain is depicted with different colors and symbols. The Abs counts represent the result of 10 simulations for H3N2 or H1N1, separately. The breadth and magnitude of Abs is considerably high in the complete experiment, between infections and after the second infection.

Time window of 20 days between infections. A window up to 6 weeks between infections (as in the principal test) manages to develop the best Abs magnitude for the first infection and therefore, a higher response magnitude for the second infection too.

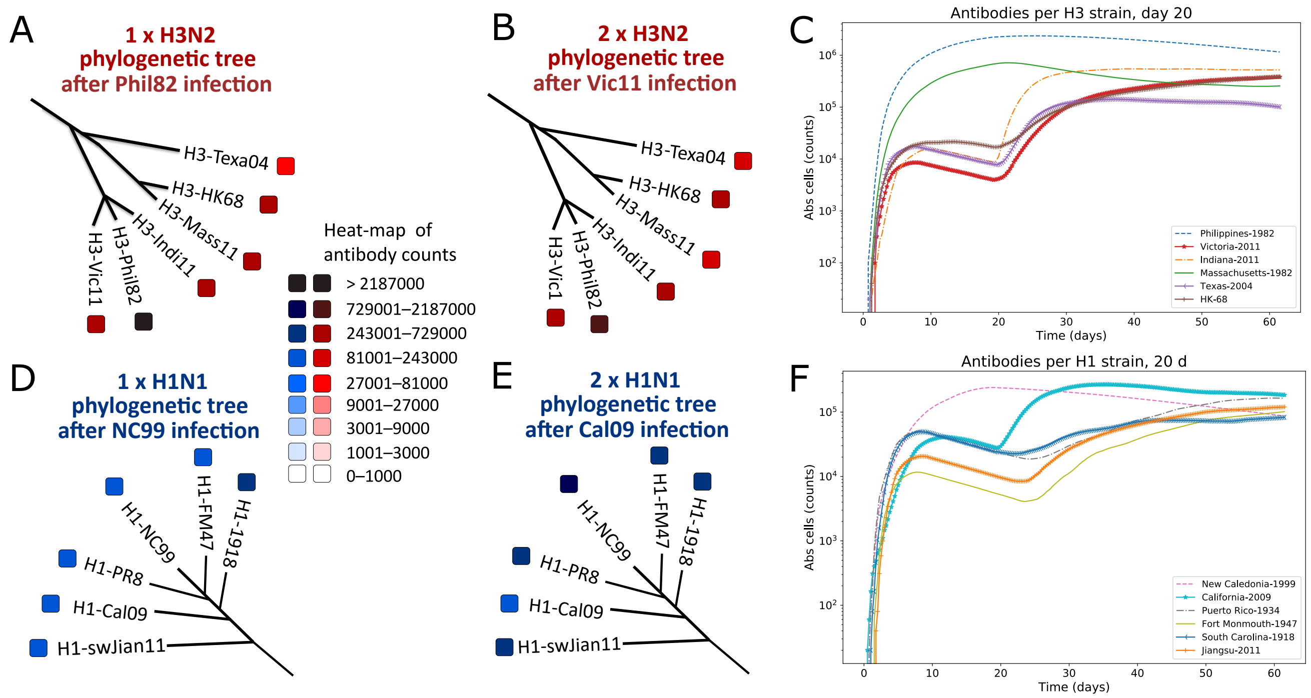


**Fig. (S11)** Test of second infection at 20 days after first infection. Test of second infection at 10 days after first infection. (A-B) A red-colored heat-map display the intervals of Abs counts for different H3N2 strains with the same characteristics of Fig. 3 of the main text. (A) Abs counts after the first influenza infection with Phil82. (B) Abs counts after the second infection with Vic11 strain. (C) The dynamics of Abs for both H3N2 infections in time with each H3 strain is depicted with different colors and symbols. (D-E) A blue-colored heat-map stand for the intervals of Abs counts for one or two consecutive H1N1 infections with the same characteristics of Fig. 3 of the main text. (D) The first H1N1 infection is with the NC99 followed by NL09 strain (E). (F) The dynamics of Abs counts for both H1N1 infections in time (days), each H1N1 strain is depicted with different colors and symbols. The Abs counts represent the result of 10 simulations for H3N2 or H1N1, separately. The breadth and magnitude of Abs is also high, as in Fig. S6, in the complete experiment, between infections and after the second infection.

Time window of 84 days between infections. A window up to 6 weeks between infections (as in the principal test) manages to develop the best Abs magnitude for the first infection and therefore, a higher response magnitude for the second infection as well. With an 84 d window, the Abs magnitude of the first infection does not support the high Abs levels for the second infection. However, the strain of infection Abs continues with great magnitude, this is due to short-term memory B-cells developed during the first infection.

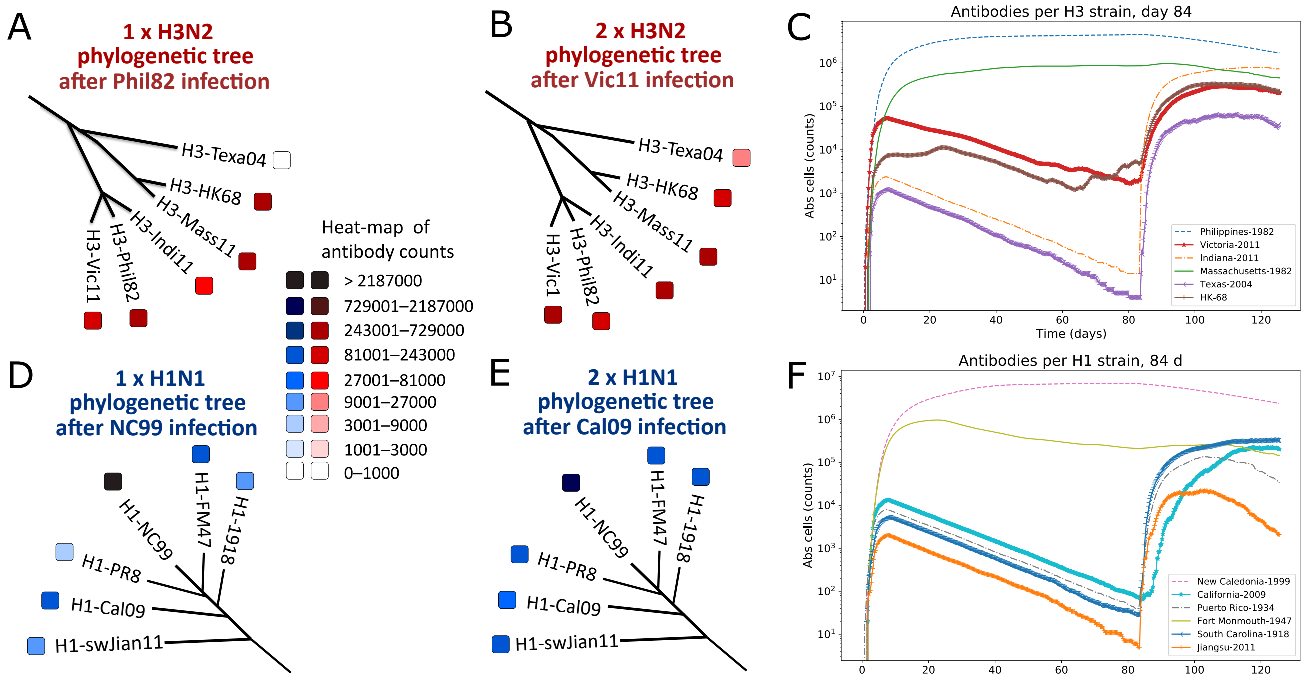


**Fig. (S12)** Test of second infection at 84 days after first infection. (A-B) A red-colored heat-map display the intervals of Abs counts for different H3N2 strains with the same characteristics of Fig. 3 of the main text. (A) Abs counts after the first influenza infection with Phil82. (B) Abs counts after the second infection with Vic11 strain. (C) The dynamics of Abs for both H3N2 infections in time with each H3 strain is depicted with different colors and symbols. (D-E) A blue-colored heat-map stand for the intervals of Abs counts for one or two consecutive H1N1 infections with the same characteristics of Fig. 3 of the main text. (D) The first H1N1 infection is with the NC99 followed by NL09 strain (E). (F) The dynamics of Abs counts for both H1N1 infections in time (days), each H1N1 strain is depicted with different colors and symbols. The Abs counts represent the result of 10 simulations for H3N2 or H1N1, separately. The breadth and magnitude of Abs is low compared to the principal test in Fig. 3 in the complete experiment.

### 8. *Commutative* tests

Fig. S13 depicts the outcome of the *Commutative* tests with inverted order of consecutive infections. The general shape of dynamics and phylogenetic trees stand for a positive commutation of infections that is dependent of the strains selection.

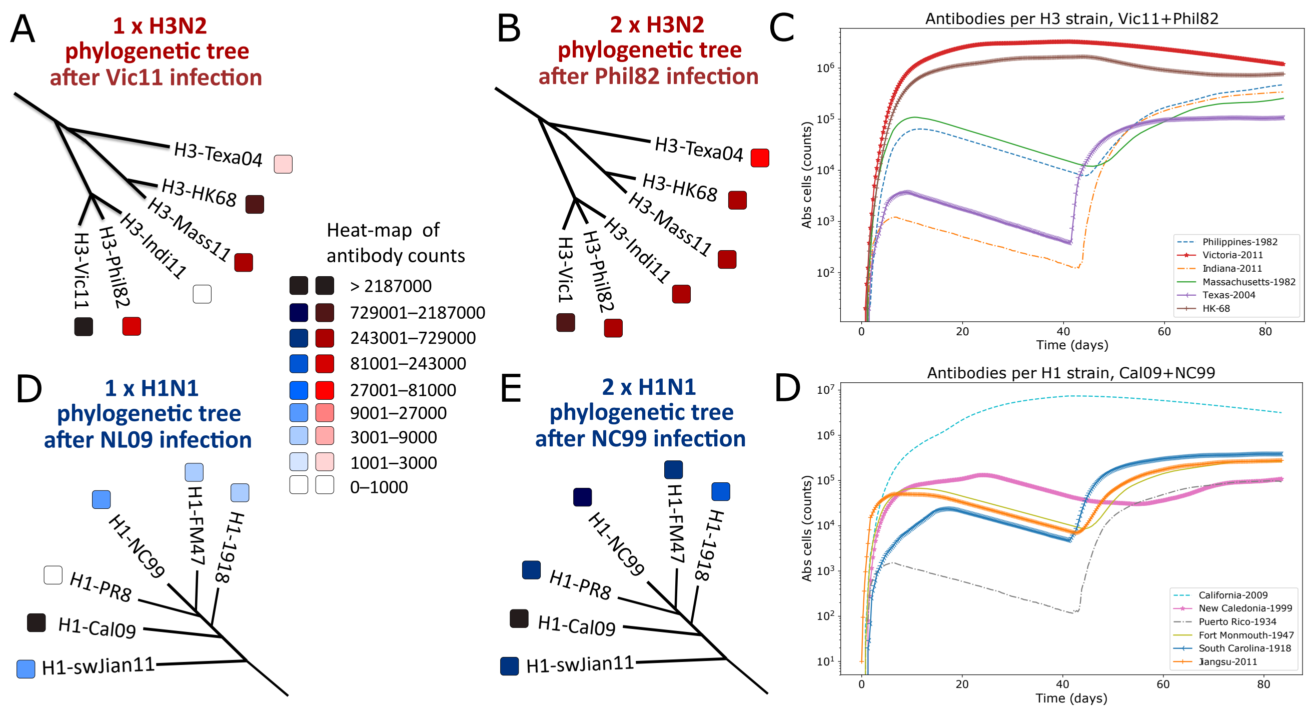


**Fig. (S13)** *Commutative* tests with inverted order of consecutive infections. (A-B) A red-colored heat-map display the intervals of Abs counts for different H3N2 strains with the same characteristics of Fig. 3 of the main text. (A) Abs counts after the first influenza infection with Vic11 strain. (B) Abs counts after the second infection with Phil82 strain. (C) The dynamics of Abs for both H3N2 infections in time with each H3 strain is depicted with different colors and symbols. (D-E) A blue-colored heat-map stand for the intervals of Abs counts for one or two consecutive H1N1 infections with the same characteristics of Fig. 3 of the main text. (D) The first H1N1 infection is with the NL09 followed by NC99 (E). (F) The dynamics of Abs counts for both H1N1 infections in time (days), each H1N1 strain is depicted with different colors and symbols. The Abs counts represent the result of 10 simulations for H3N2 or H1N1, separately. The general shape of dynamics and phylogenetic trees stand for a positive commutation of infections, at least for the tested ones.

### 9. Simultaneous infection test

We explore the simultaneous infection scenario using the same influenza strains of the main test for H3N2 and H1N1, separately. For influenza H3N2, we use Phill82+Vic11. For influenza H1N1, we use NC99+Cal09. Fig. (S14) shows the results of this test.

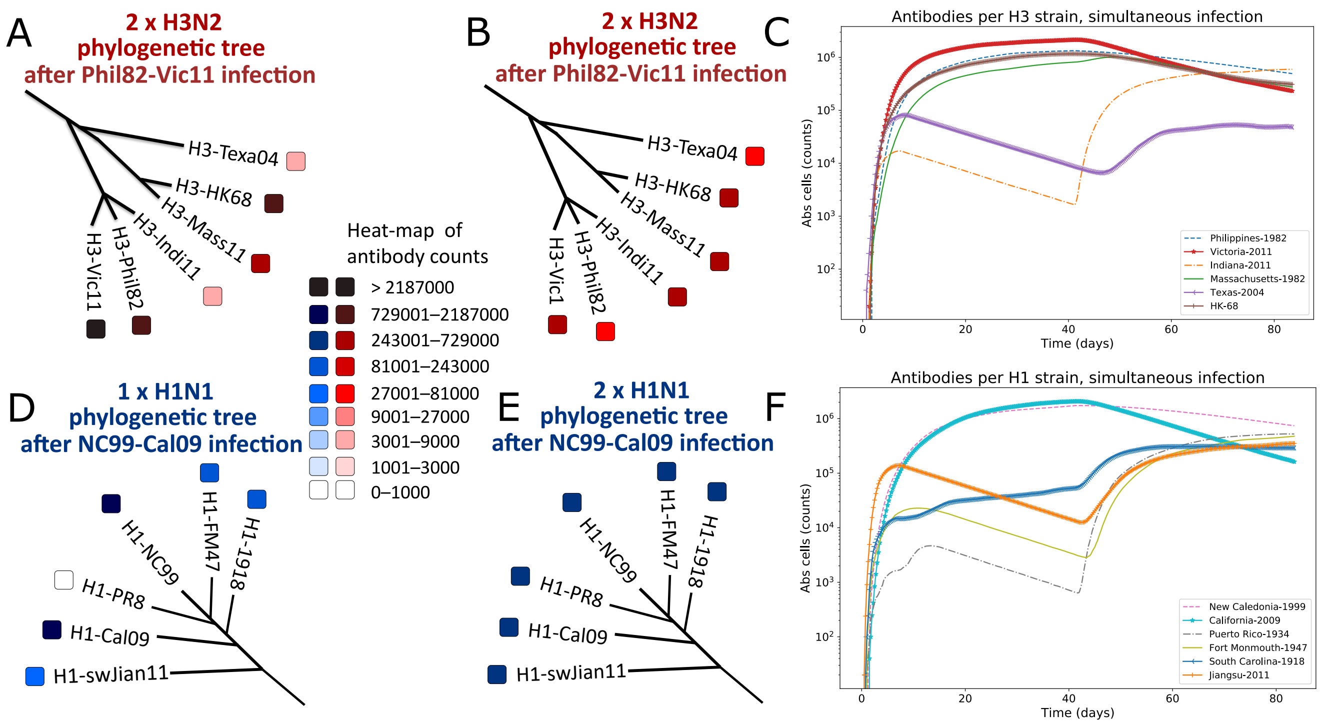


**Fig. (S14)** Simultaneous infection test. (A-B) A red-colored heat-map display the intervals of Abs counts for different H3N2 strains with the same characteristics of Fig. 3 of the main text. (A) Abs counts after the first influenza infection with Phil82 and Vic11. (B) Abs counts after the second and equivalent infection. (C) The dynamics of Abs for both H3N2 infections in time with each H3 strain is depicted with different colors and symbols. (D-E) A blue-colored heat-map stand for the intervals of Abs counts for one or two consecutive H1N1 infections with the same characteristics of Fig. 3 of the main text. (D) The first H1N1 infection is with the NC99-Cal09 followed by equal scheme of infection (E). (F) The dynamics of Abs counts for both H1N1 infections in time (days), each H1N1 strain is depicted with different colors and symbols. The Abs counts represent the result of 10 simulations for H3N2 or H1N1, separately. The second infection depicts similar results to the main test and the first infection produces higher Abs counts compared to the main test. This may have implications of cocktails of antigens for vaccine designs.

### 10. Principal Test simulations of H3N2 strain

Simulation 1 of the Principal Test with H3N2 strain.

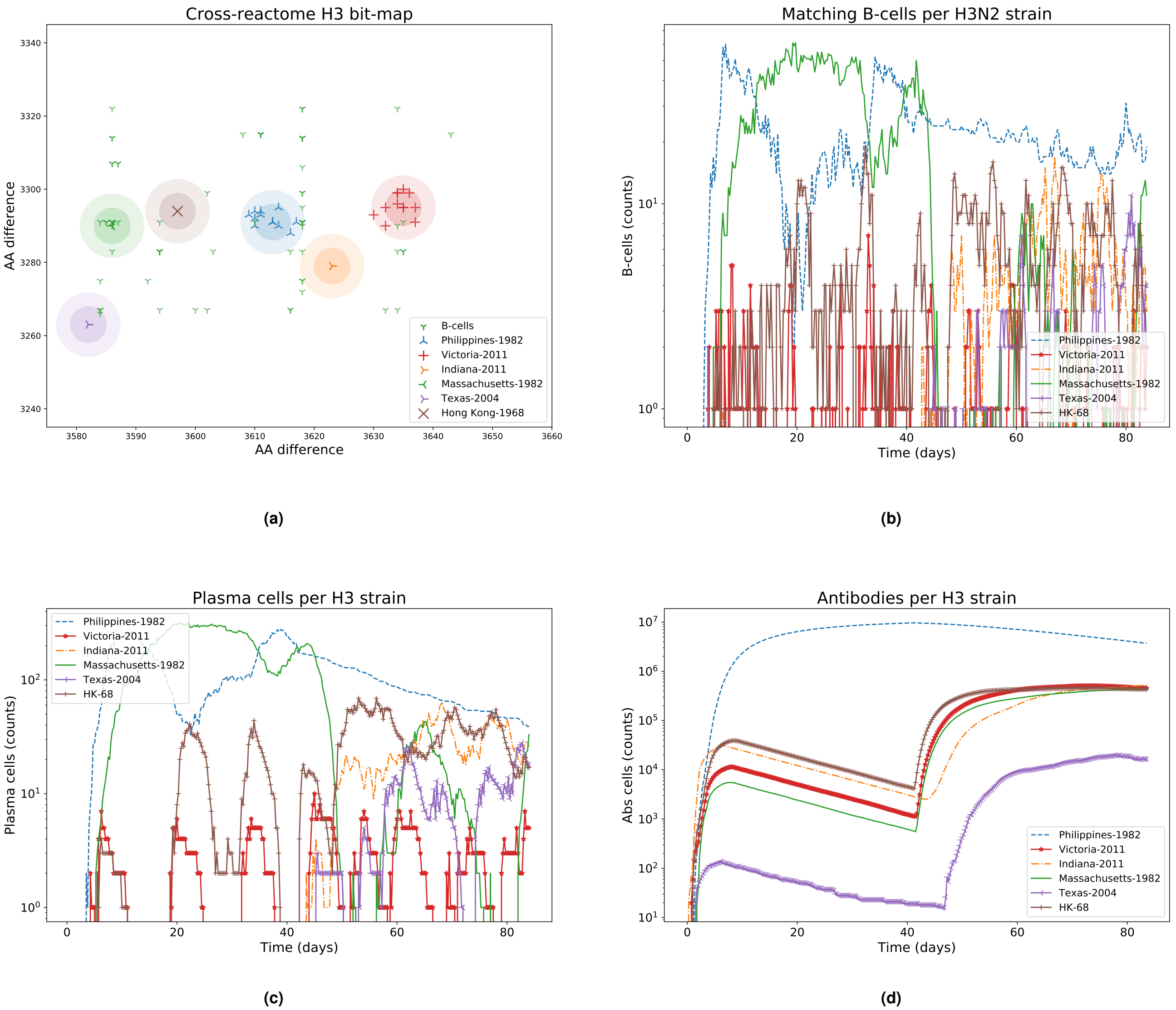


**Fig. (S15)** H3N2 Influenza strains *shape-space* and immune system components dynamics, Test 1. (A) B-cell population covering most of the H3N2 influenza strains clusters after the Vic11 infection. (B) A B-cell that falls into the diameter of a strain cluster is considered as a matching B-cell for that strain. The quantity of B-cells inside each cluster is plotted respect to time. (C) A high-affinity B-cell has the chance to generate a plasma cell which decay according to its half-life. Due to a combined effect of the high threshold for the principal area during the second infection and AM, the plasma cells in the Vic11 infection manage to remain more time than for the Phil82 infection. (D) The antibody population for each strain is generated by plasma cells targeting that corresponding strain.

Simulation 2 of the Principal Test with H3N2 strain.

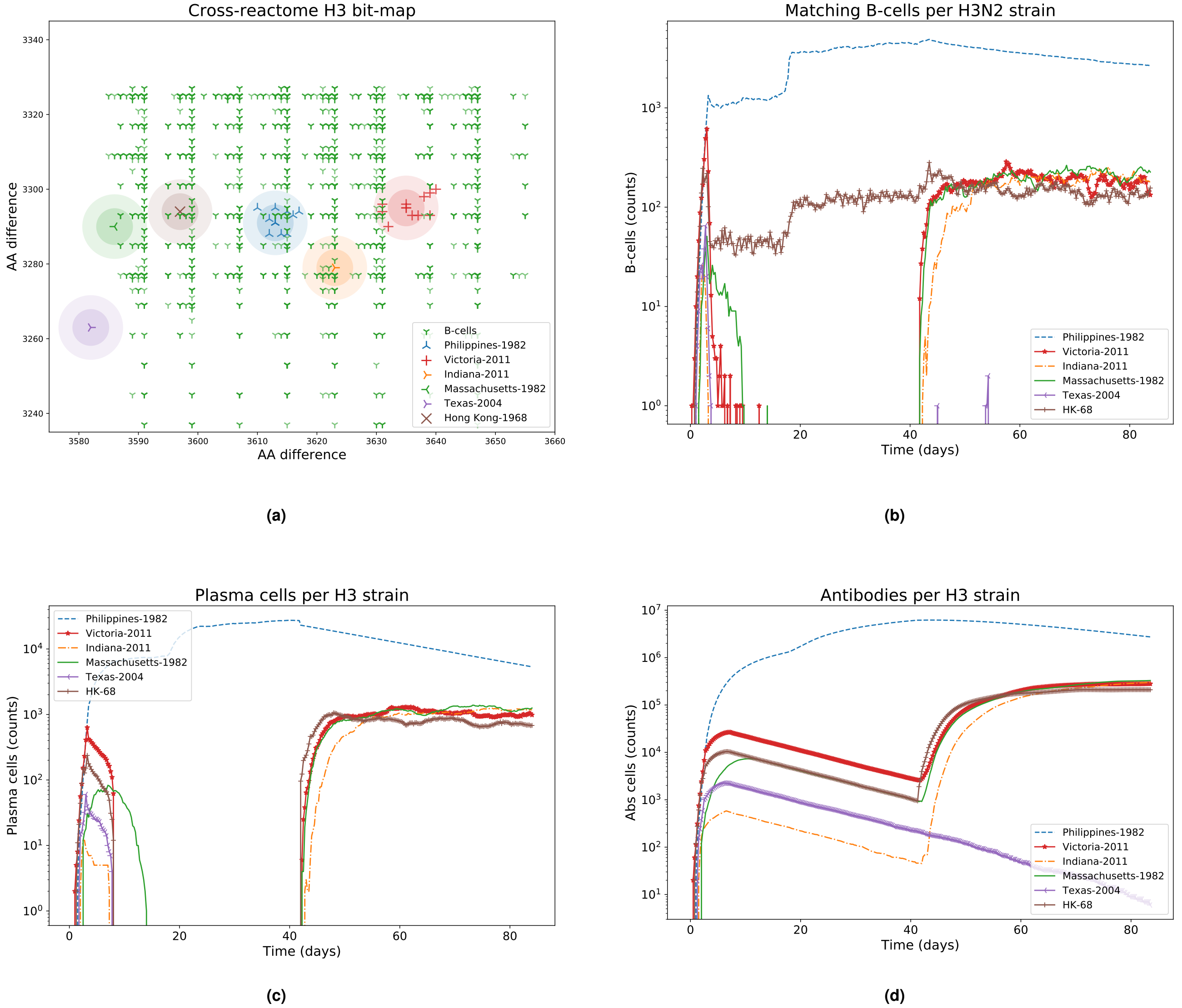


**Fig. (S16)** H3N2 Influenza strains *shape-space* and immune system components dynamics, Test 2.

Simulation 3 of the Principal Test with H3N2 strain.

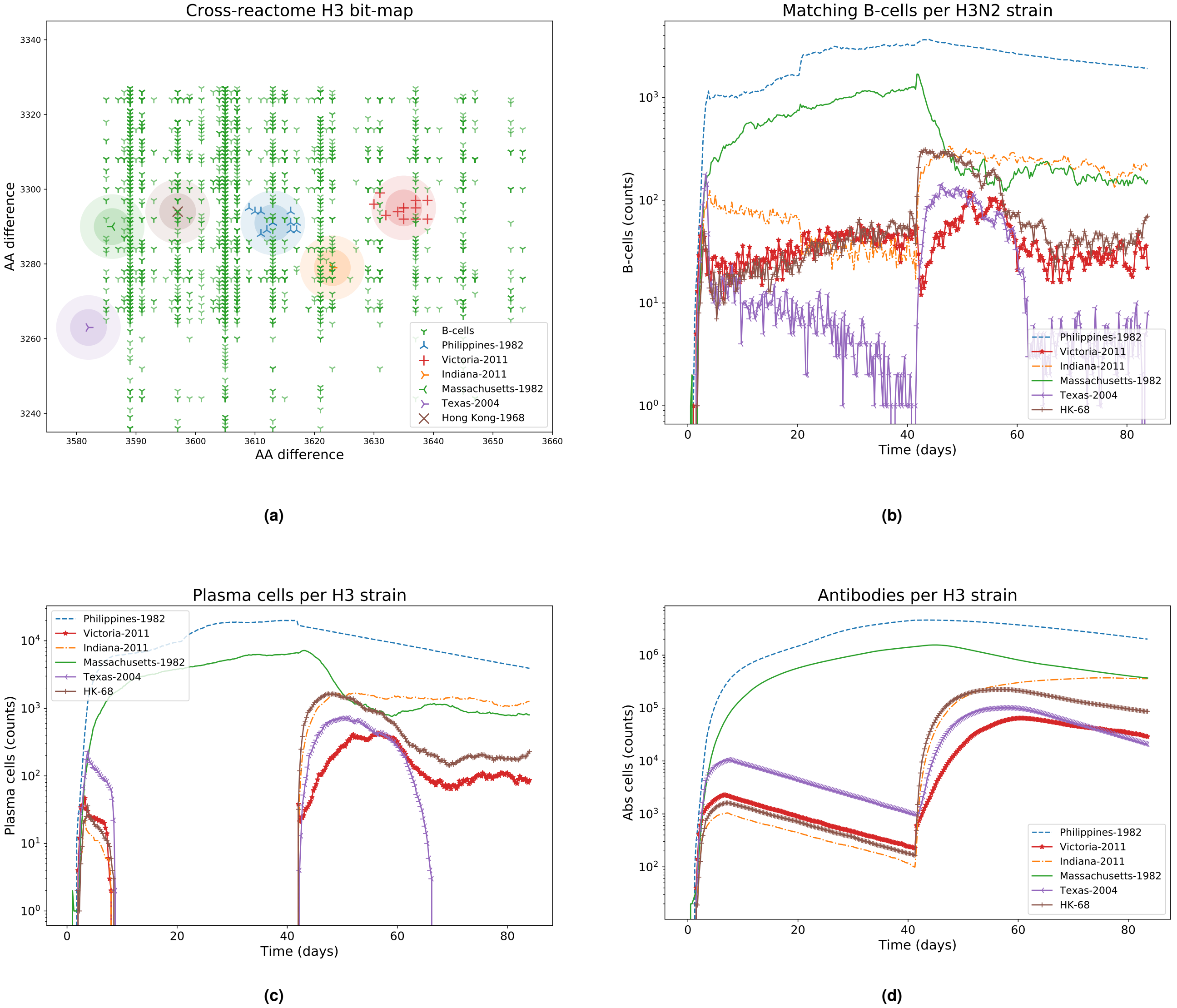


**Fig. (S17)** H3N2 Influenza strains *shape-space* and immune system components dynamics, Test 3.

Simulation 4 of the Principal Test with H3N2 strain.

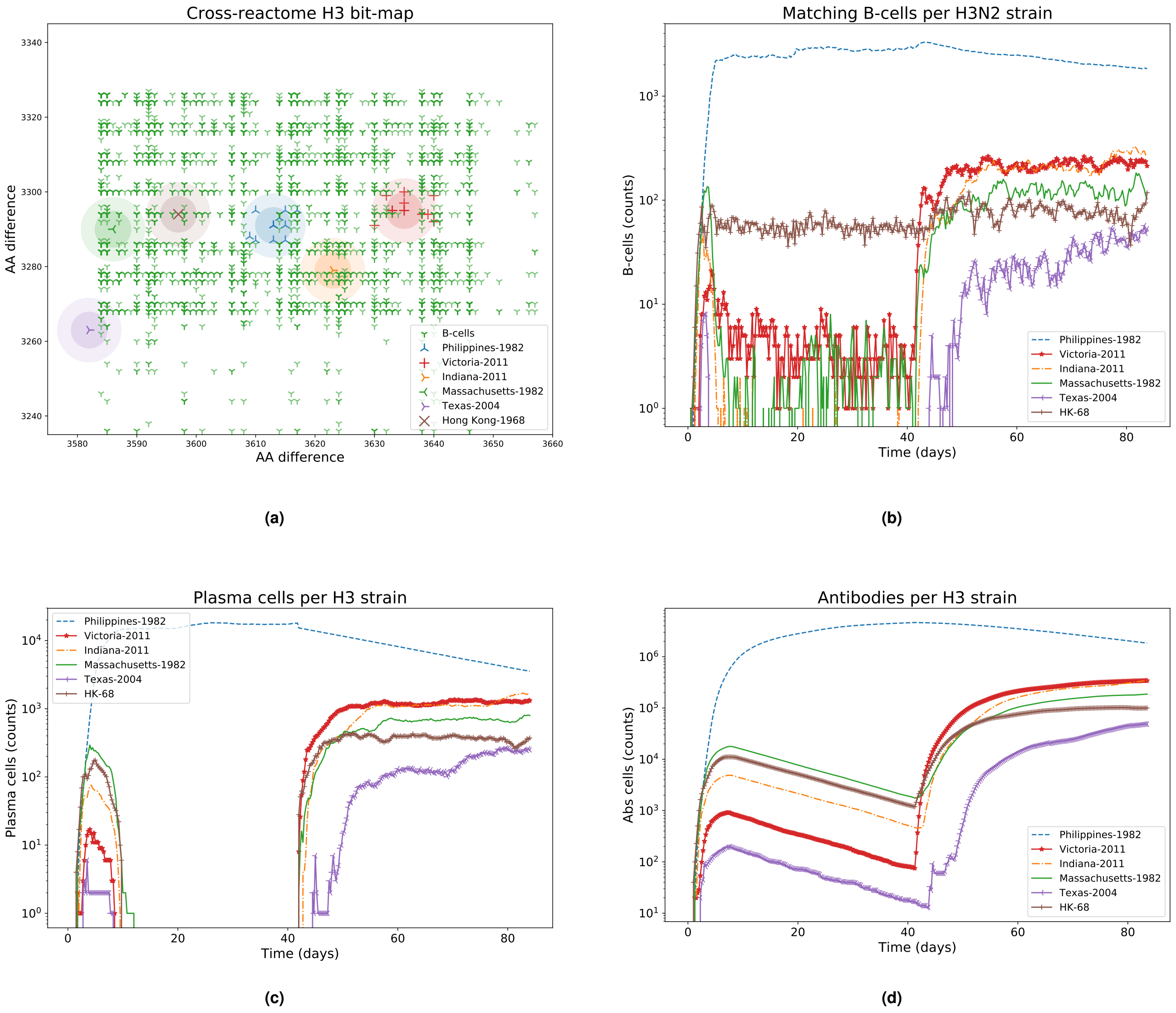


**Fig. (S18)** H3N2 Influenza strains *shape-space* and immune system components dynamics, Test 4.

Simulation 5 of the Principal Test with H3N2 strain.

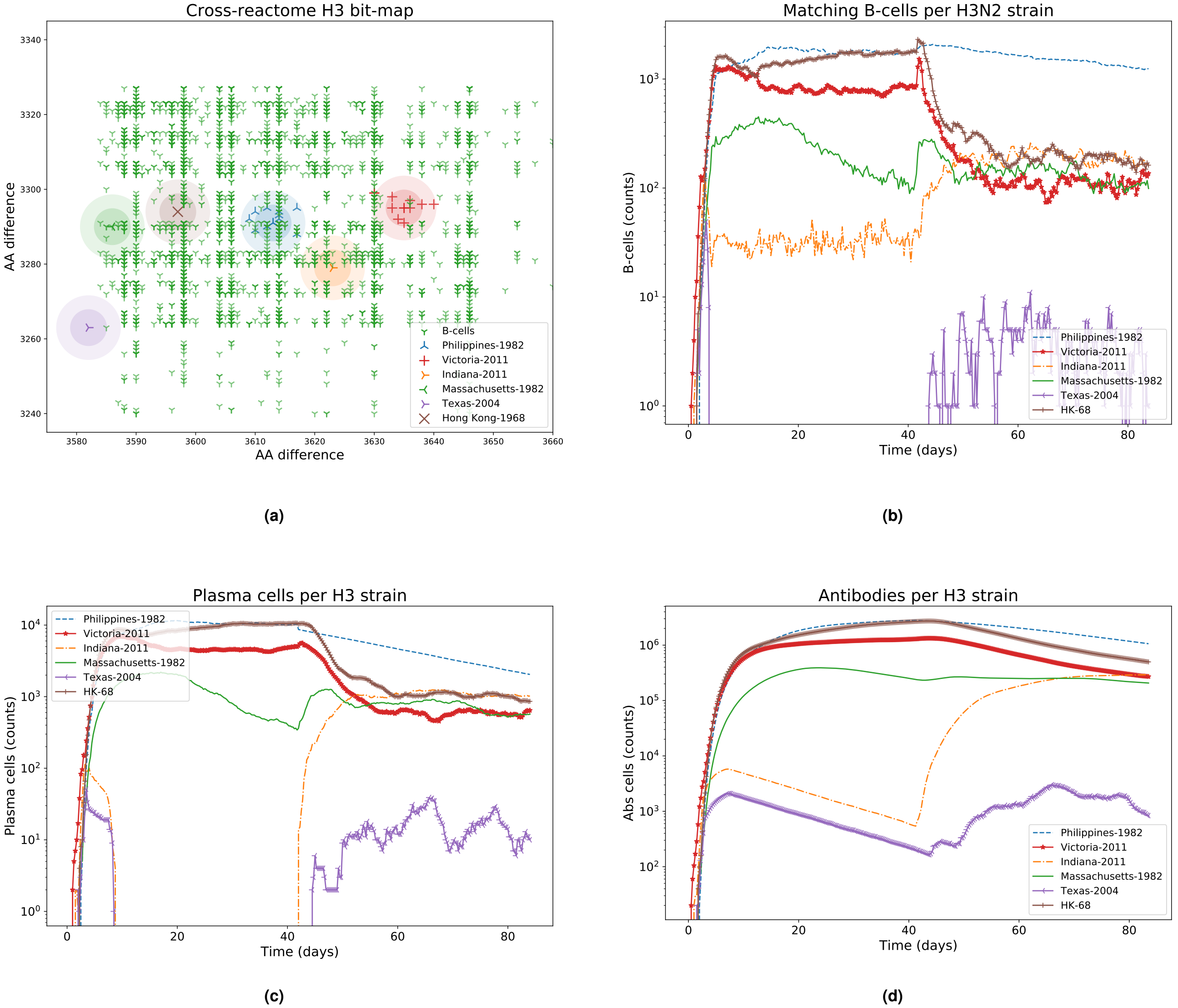


**Fig. (S19)** H3N2 Influenza strains *shape-space* and immune system components dynamics, Test 5.

Simulation 6 of the Principal Test with H3N2 strain.

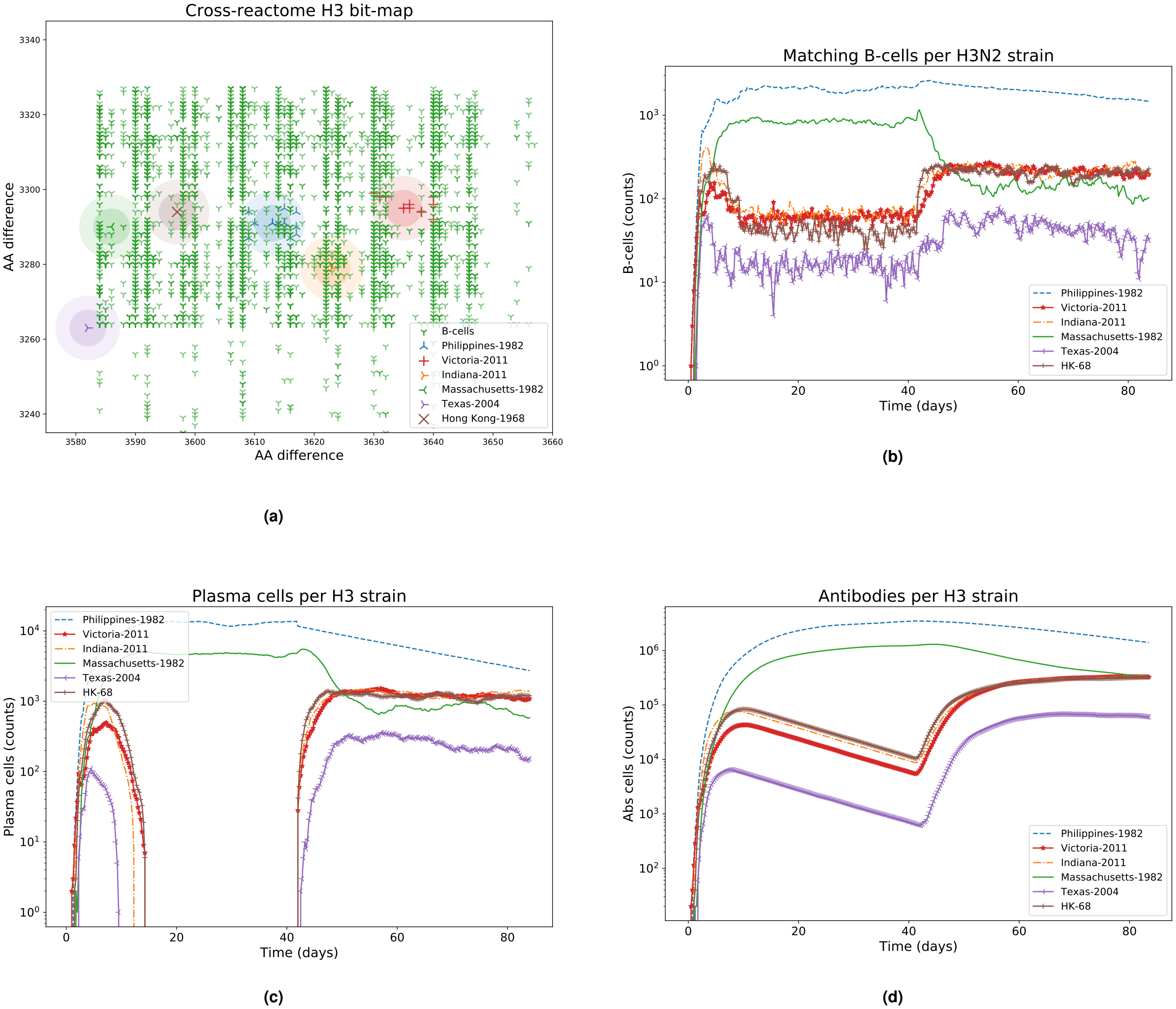


**Fig. (S20)** H3N2 Influenza strains *shape-space* and immune system components dynamics, Test 6.

Simulation 7 of the Principal Test with H3N2 strain.

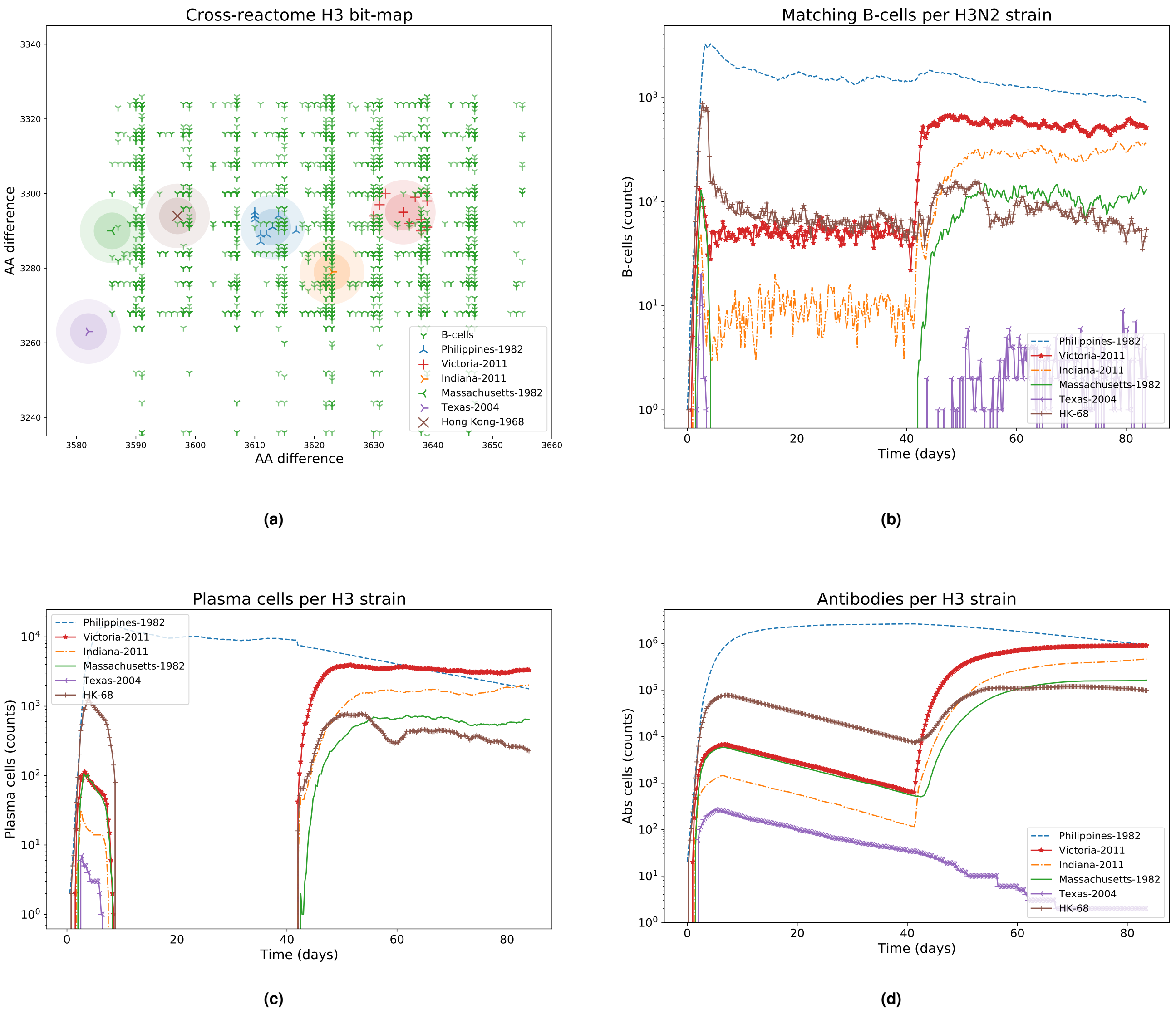


**Fig. (S21)** H3N2 Influenza strains *shape-space* and immune system components dynamics, Test 7.

Simulation 8 of the Principal Test with H3N2 strain.

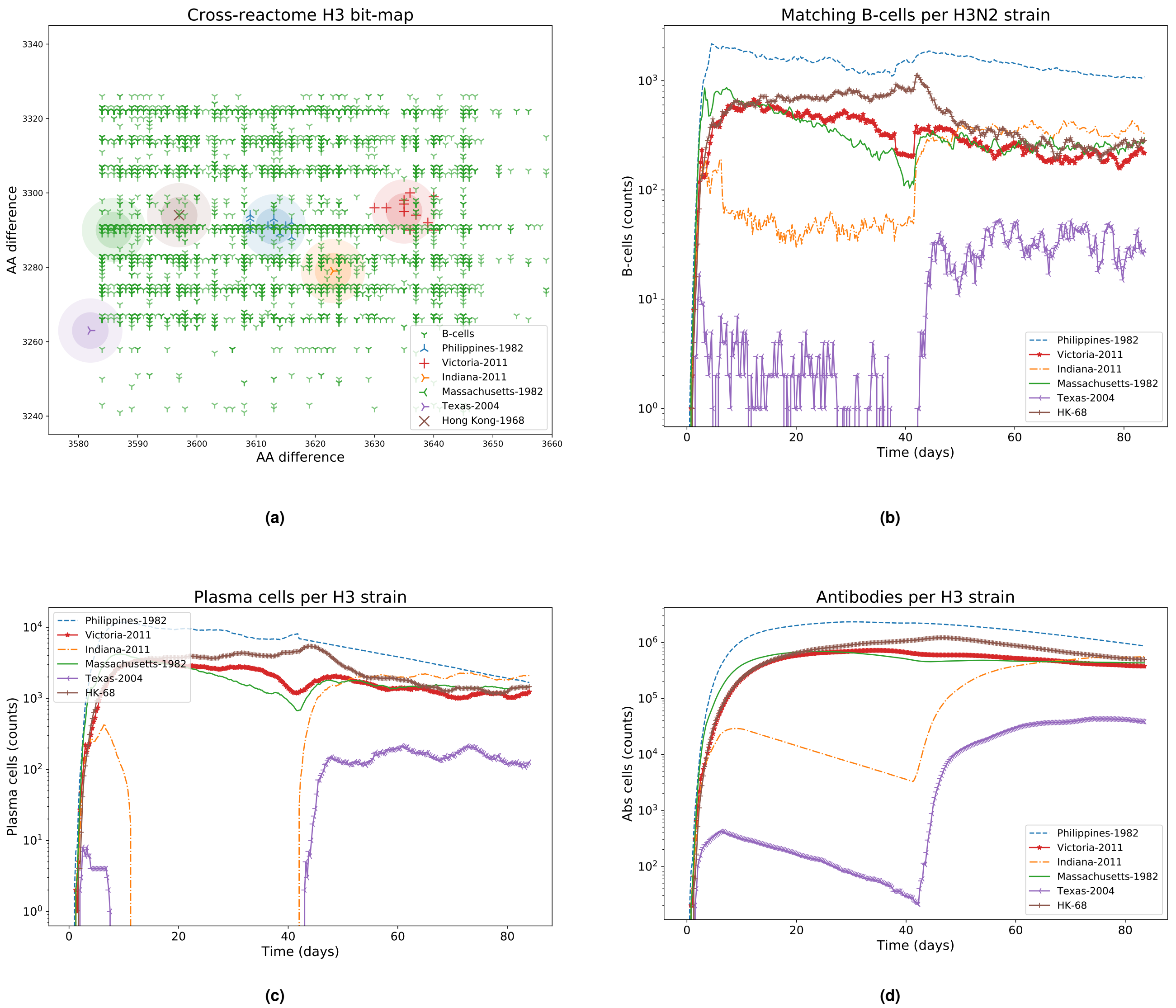


**Fig. (S22)** H3N2 Influenza strains *shape-space* and immune system components dynamics, Test 8.

Simulation 9 of the Principal Test with H3N2 strain.

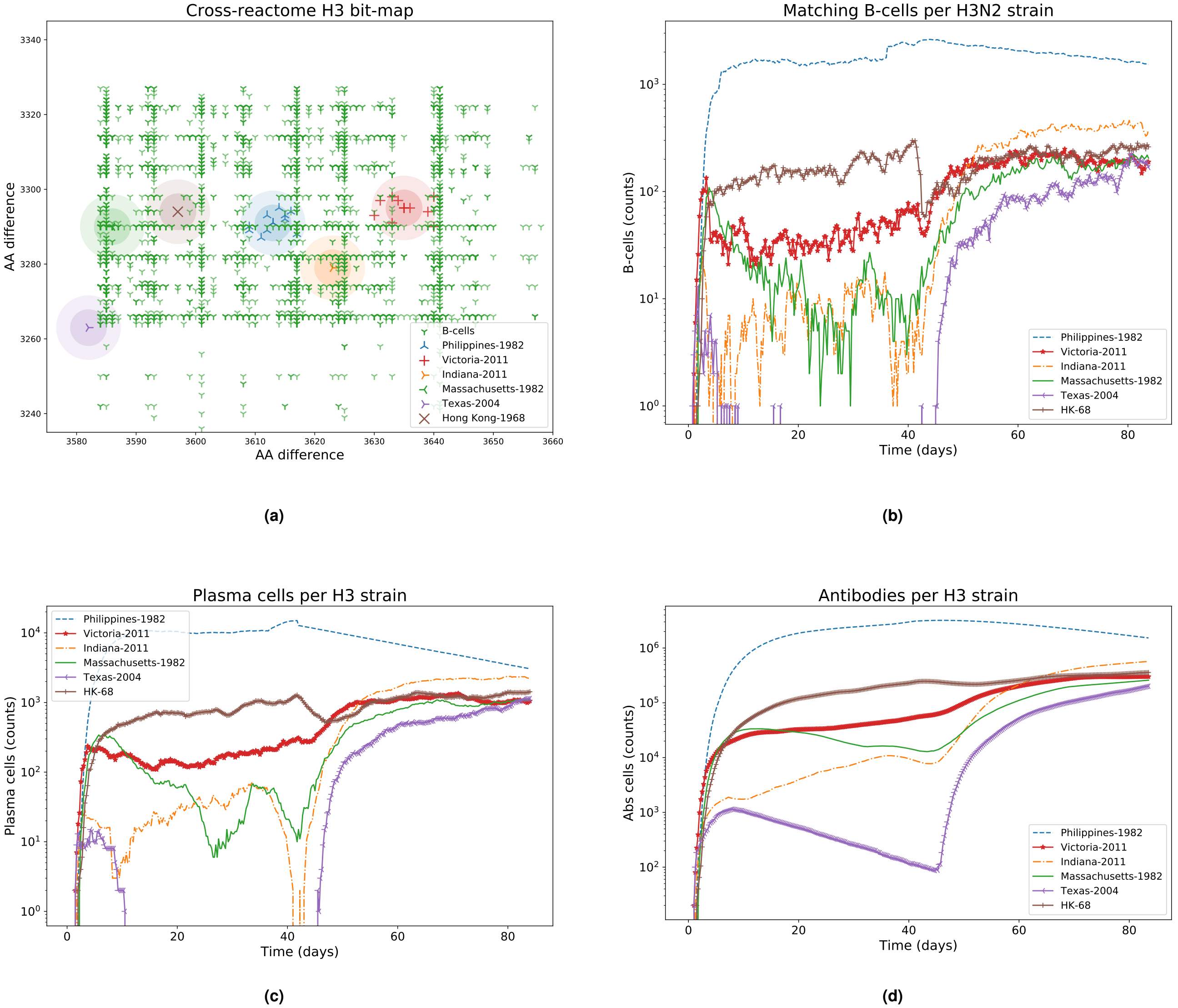


**Fig. (S23)** H3N2 Influenza strains *shape-space* and immune system components dynamics, Test 9.

### Simulation 10 of the Principal Test with H3N2 strain.

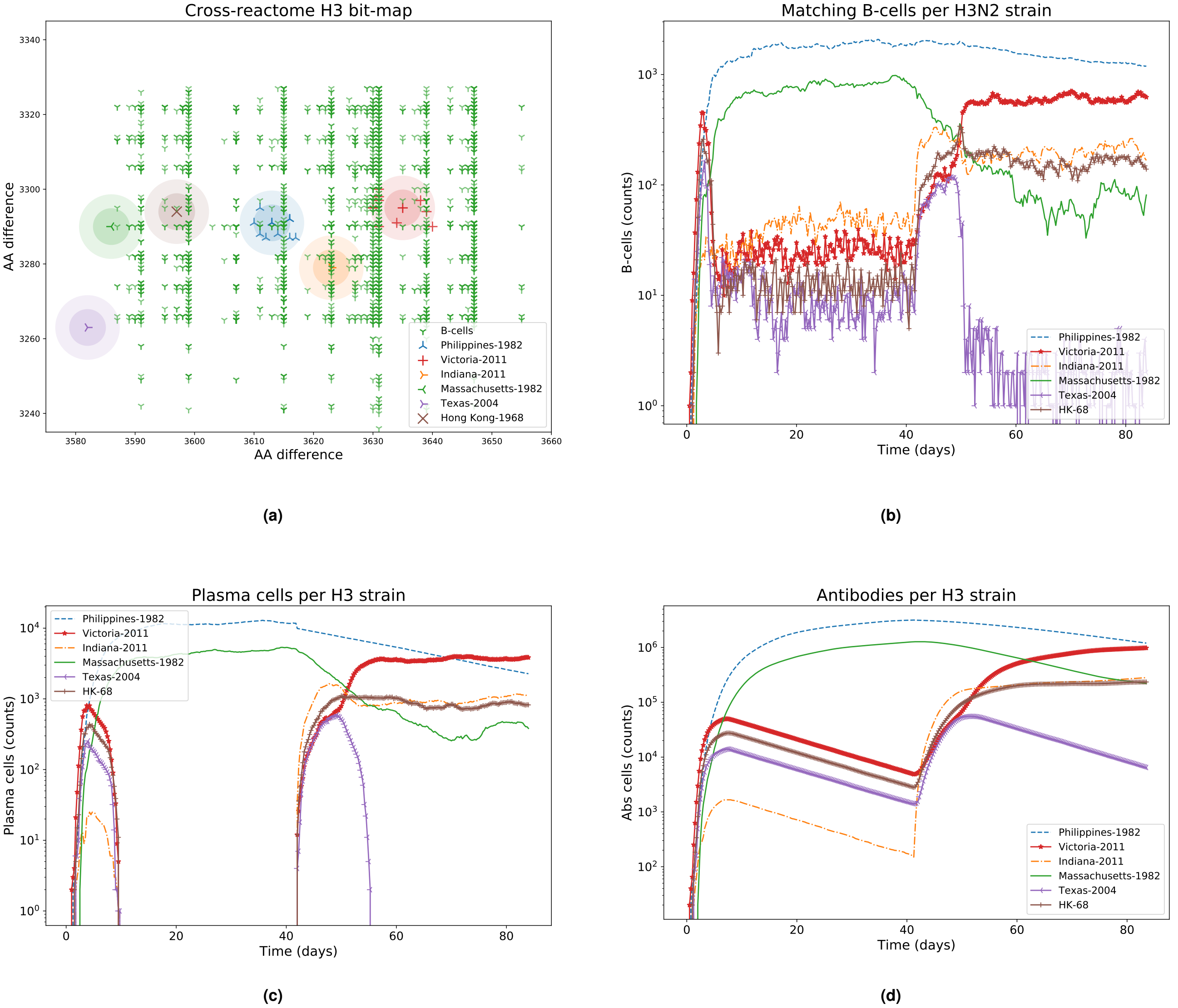


**Fig. (S24)** H3N2 Influenza strains *shape-space* and immune system components dynamics, Test 10.

### 11. Principal Test simulations of H1N1 strain

Simulation 1 of the Principal Test with H1N1 strain.

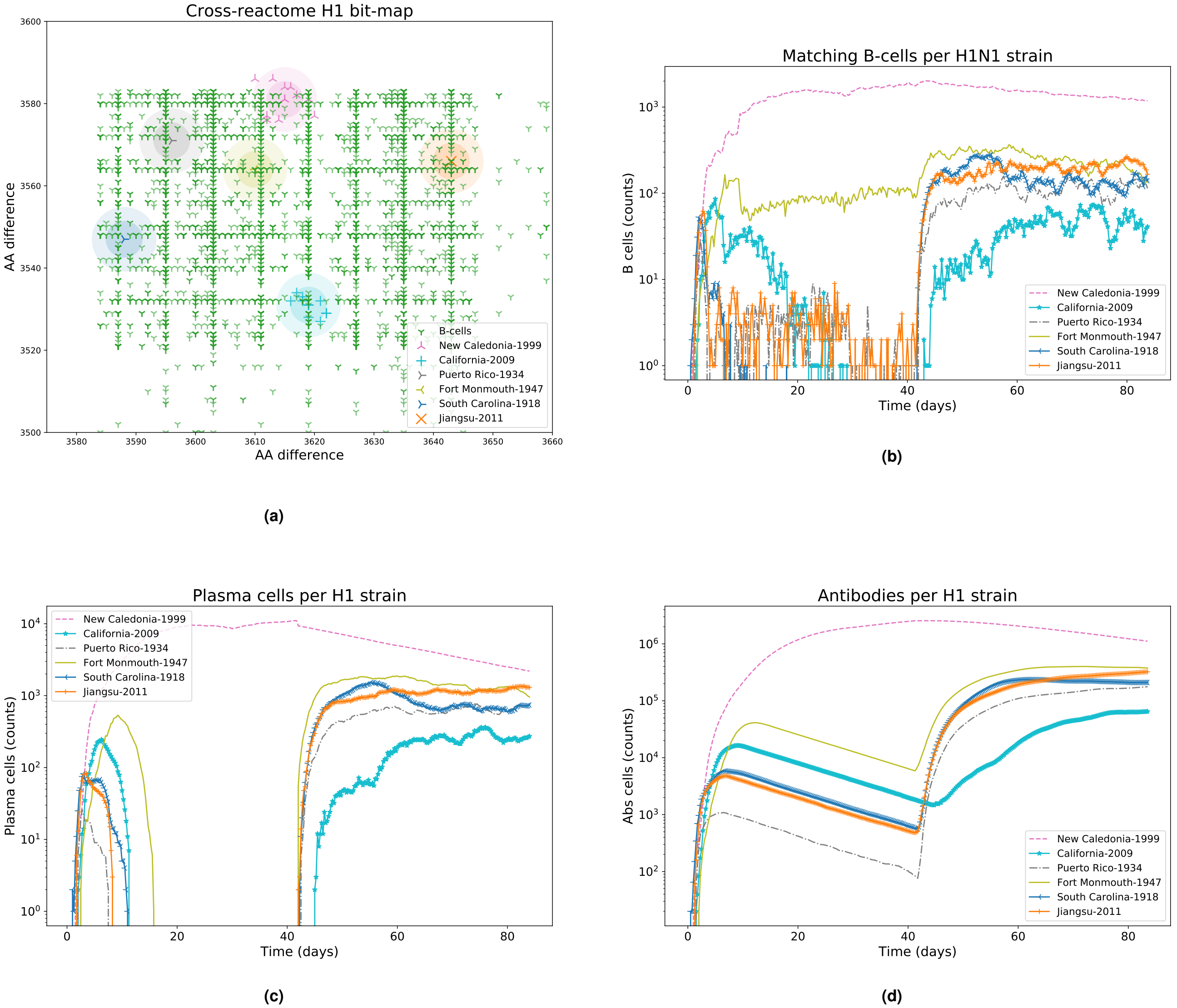


**Fig. (S25)** H1N1 Influenza strains *shape-space* and immune system components dynamics, Test 1. (A) B-cell population covering most of the H1N1 influenza strains clusters after the Cal09 infection. (B) A B-cell that falls into the diameter of a strain cluster is considered as a matching B-cell for that strain. The quantity of B-cells inside each cluster is plotted respect to time. (C) A high-affinity B-cell has the chance to generate a plasma cell which decay according to its half-life. Due to a combined effect of the high threshold for the principal area during the second infection and AM, the plasma cells in the NC99 infection manage to remain more time than for the Cal09 infection. (D) The antibody population for each strain is generated by plasma cells targeting that corresponding strain.

Simulation 2 of the Principal Test with H1N1 strain.

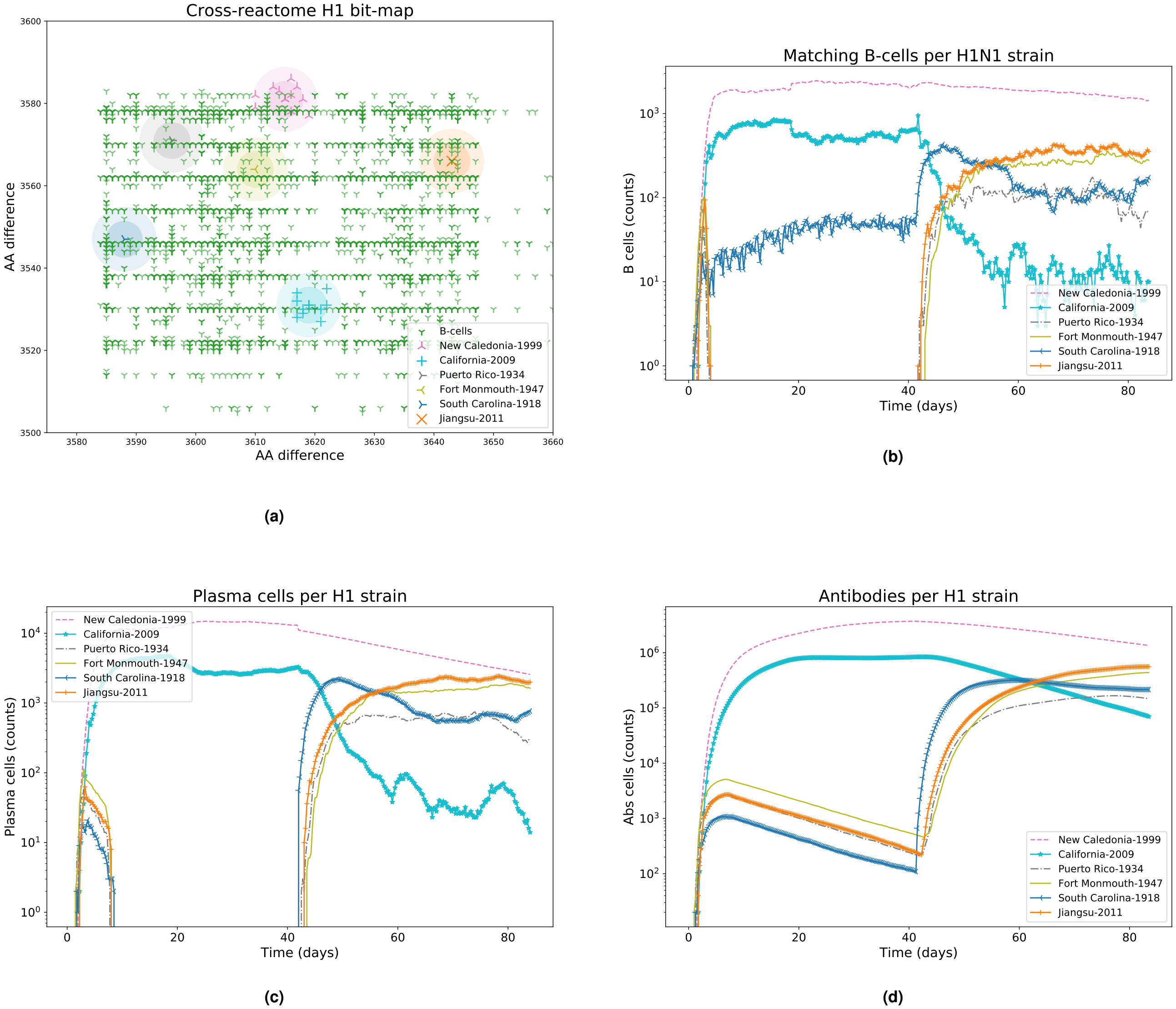


**Fig. (S26)** H1N1 Influenza strains *shape-space* and immune system components dynamics, Test 2.

Simulation 3 of the Principal Test with H1N1 strain.

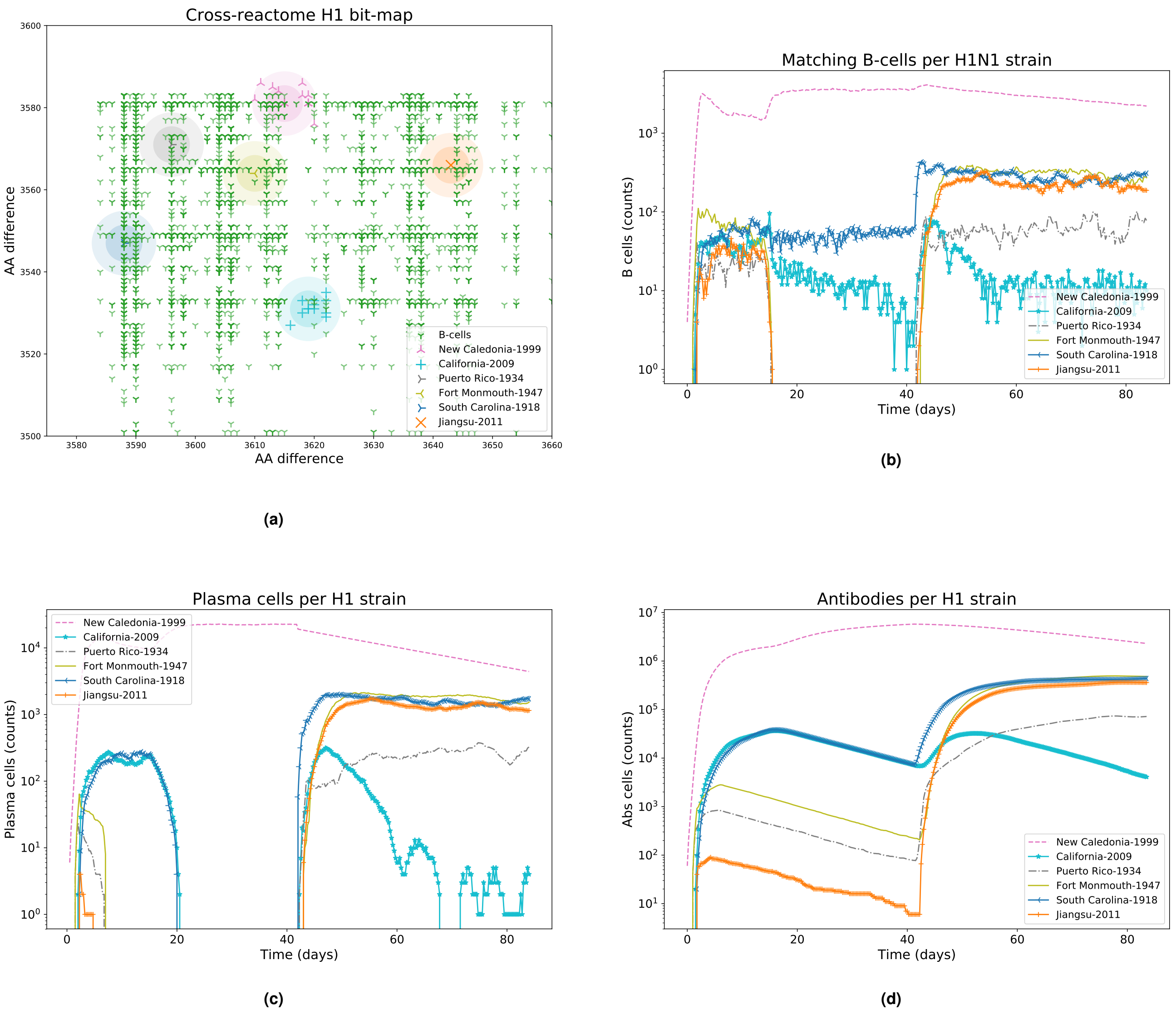


**Fig. (S27)** H1N1 Influenza strains *shape-space* and immune system components dynamics, Test 3.

Simulation 4 of the Principal Test with H1N1 strain.

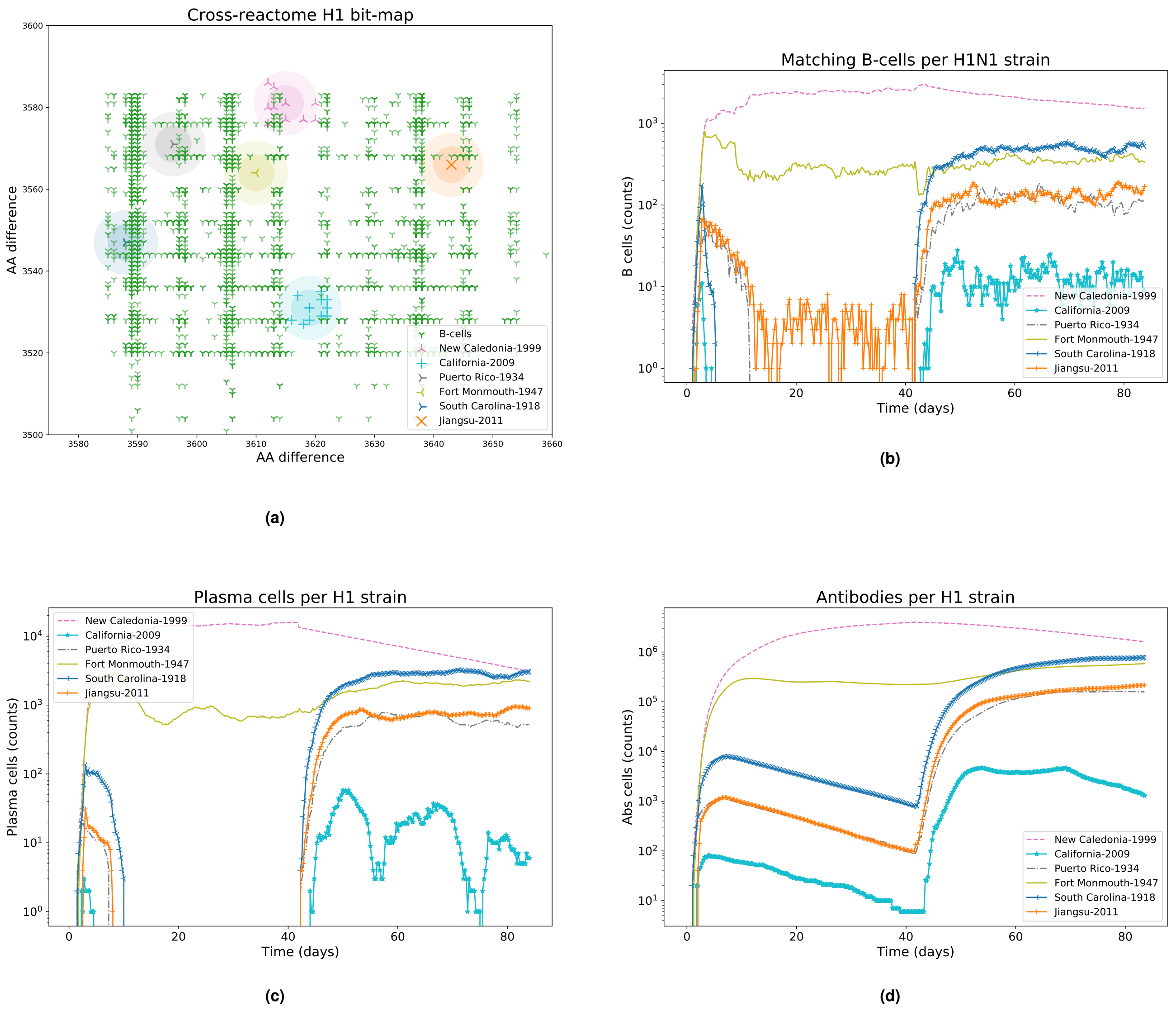


**Fig. (S28)** H1N1 Influenza strains *shape-space* and immune system components dynamics, Test 4.

Simulation 5 of the Principal Test with H1N1 strain.

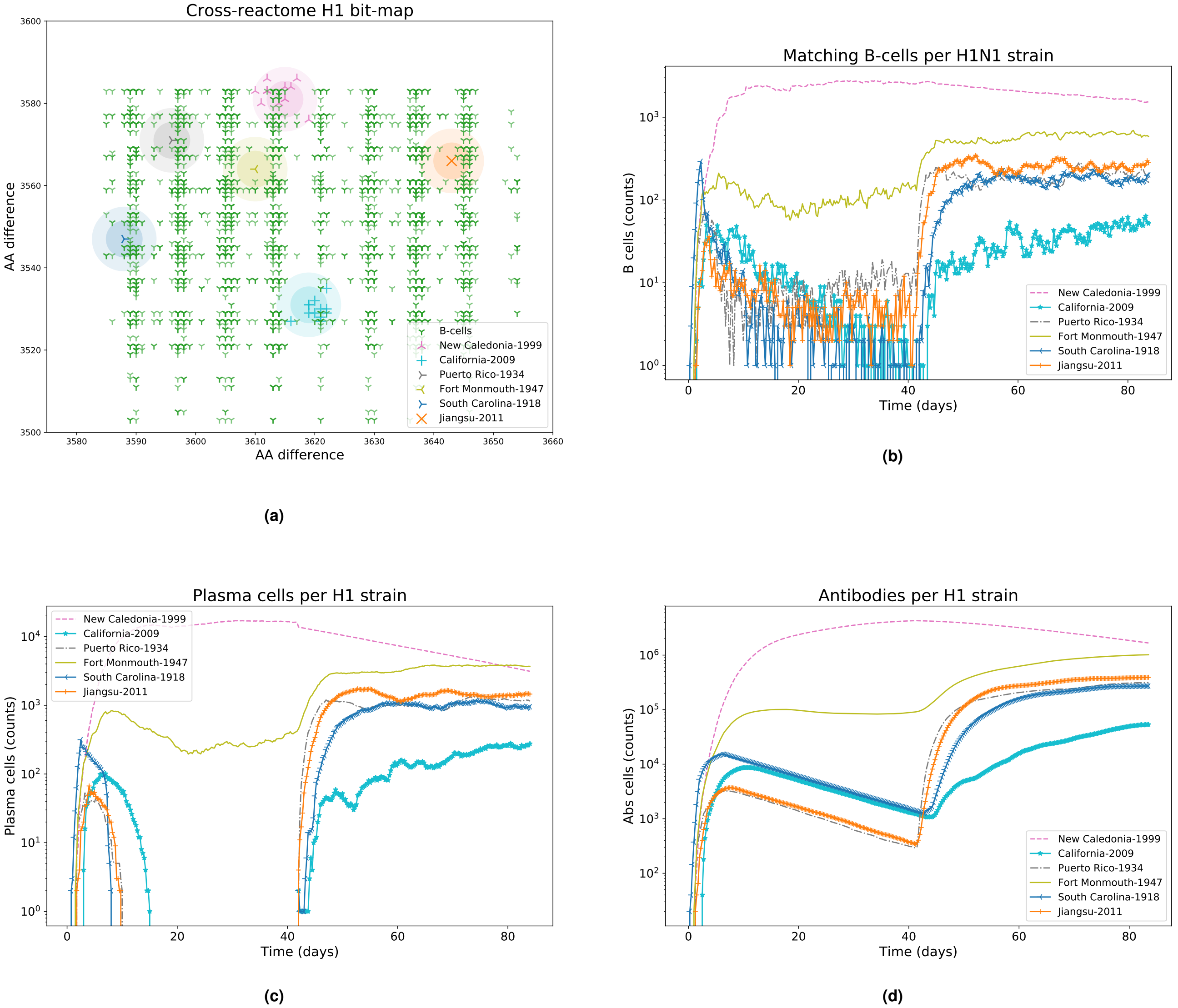


**Fig. (S29)** H1N1 Influenza strains *shape-space* and immune system components dynamics, Test 5.

Simulation 6 of the Principal Test with H1N1 strain.

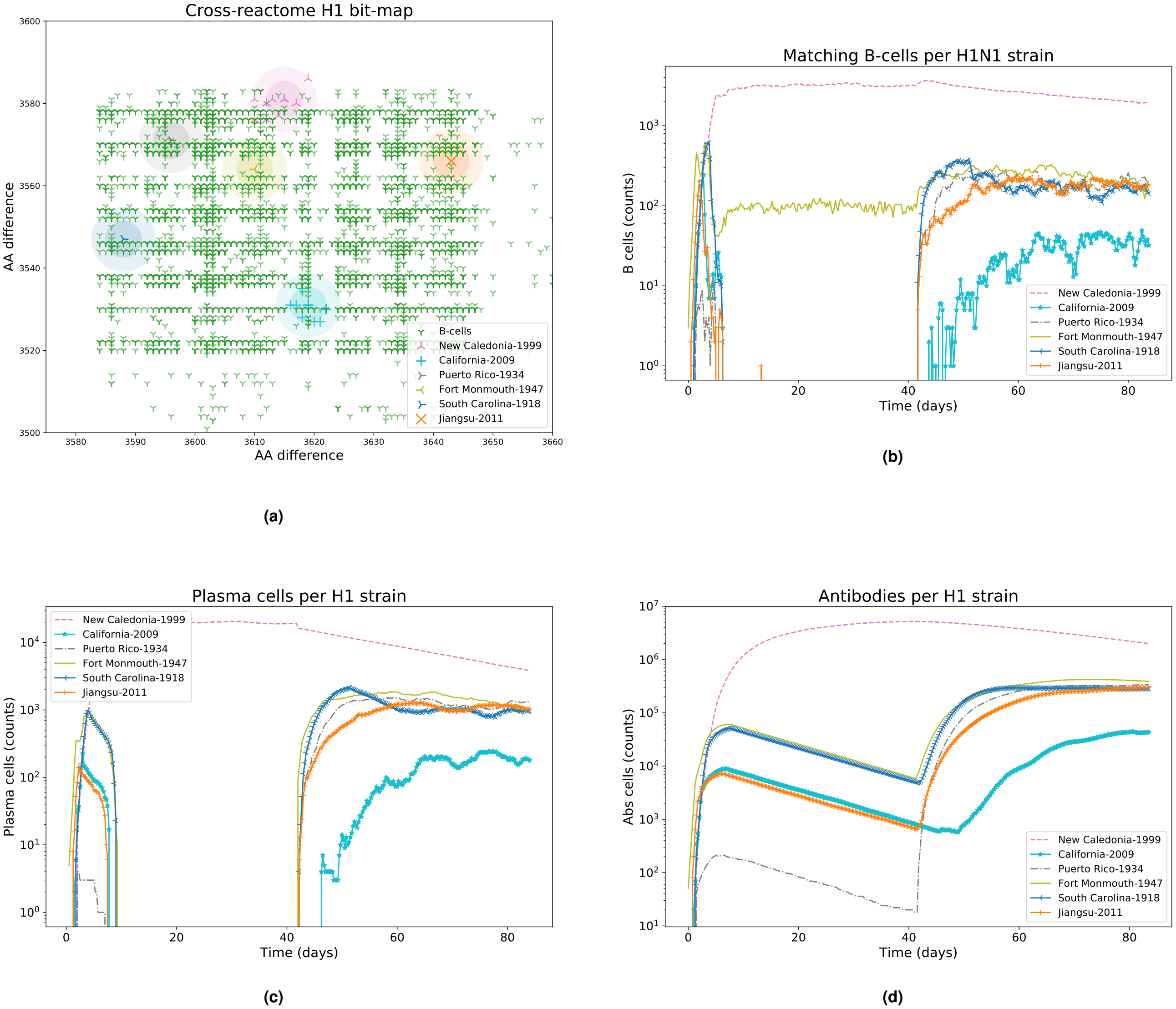


**Fig. (S30)** H1N1 Influenza strains *shape-space* and immune system components dynamics, Test 6.

Simulation 7 of the Principal Test with H1N1 strain.

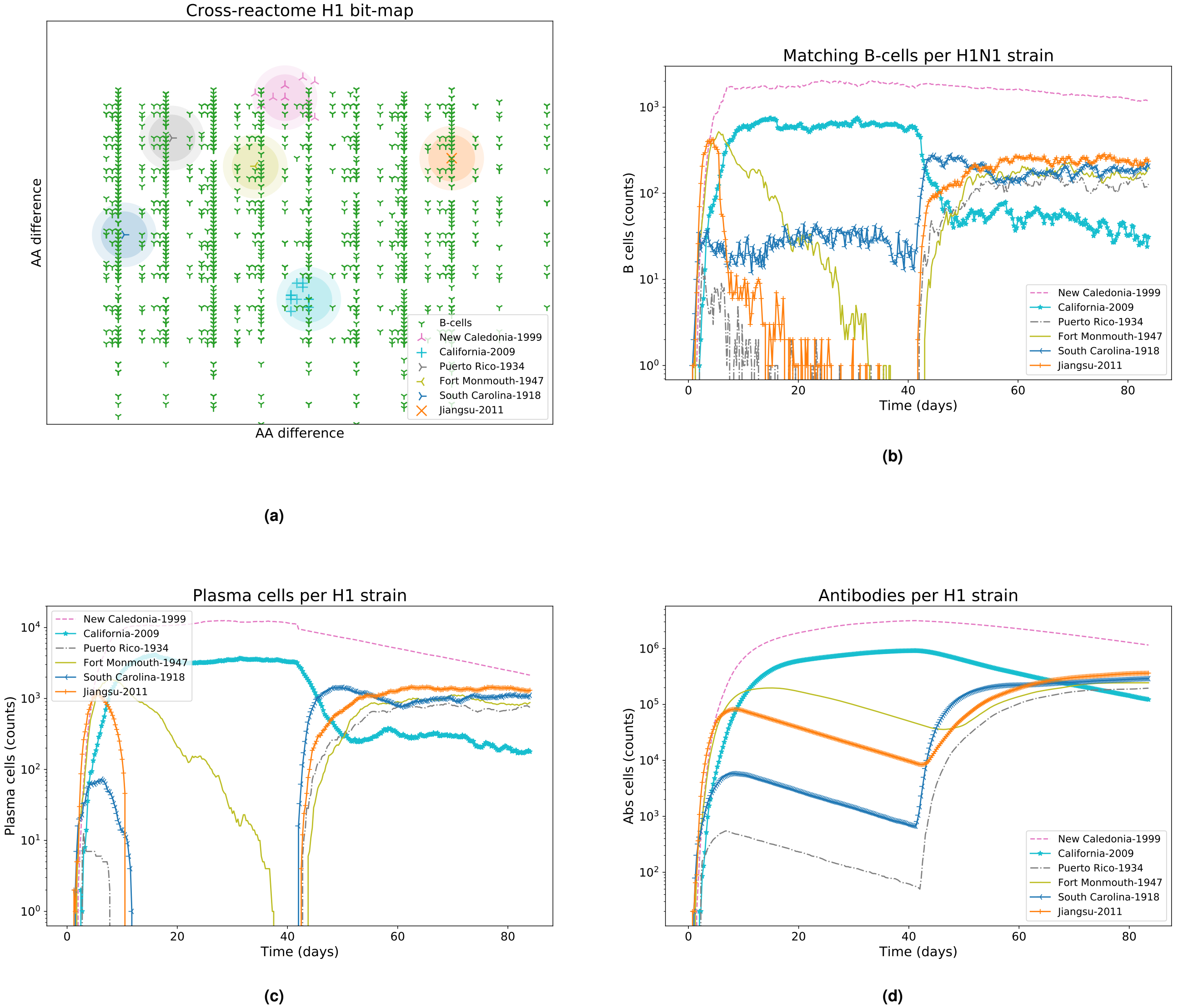


**Fig. (S31)** H1N1 Influenza strains *shape-space* and immune system components dynamics, Test 7.

 

Simulation 8 of the Principal Test with H1N1 strain.

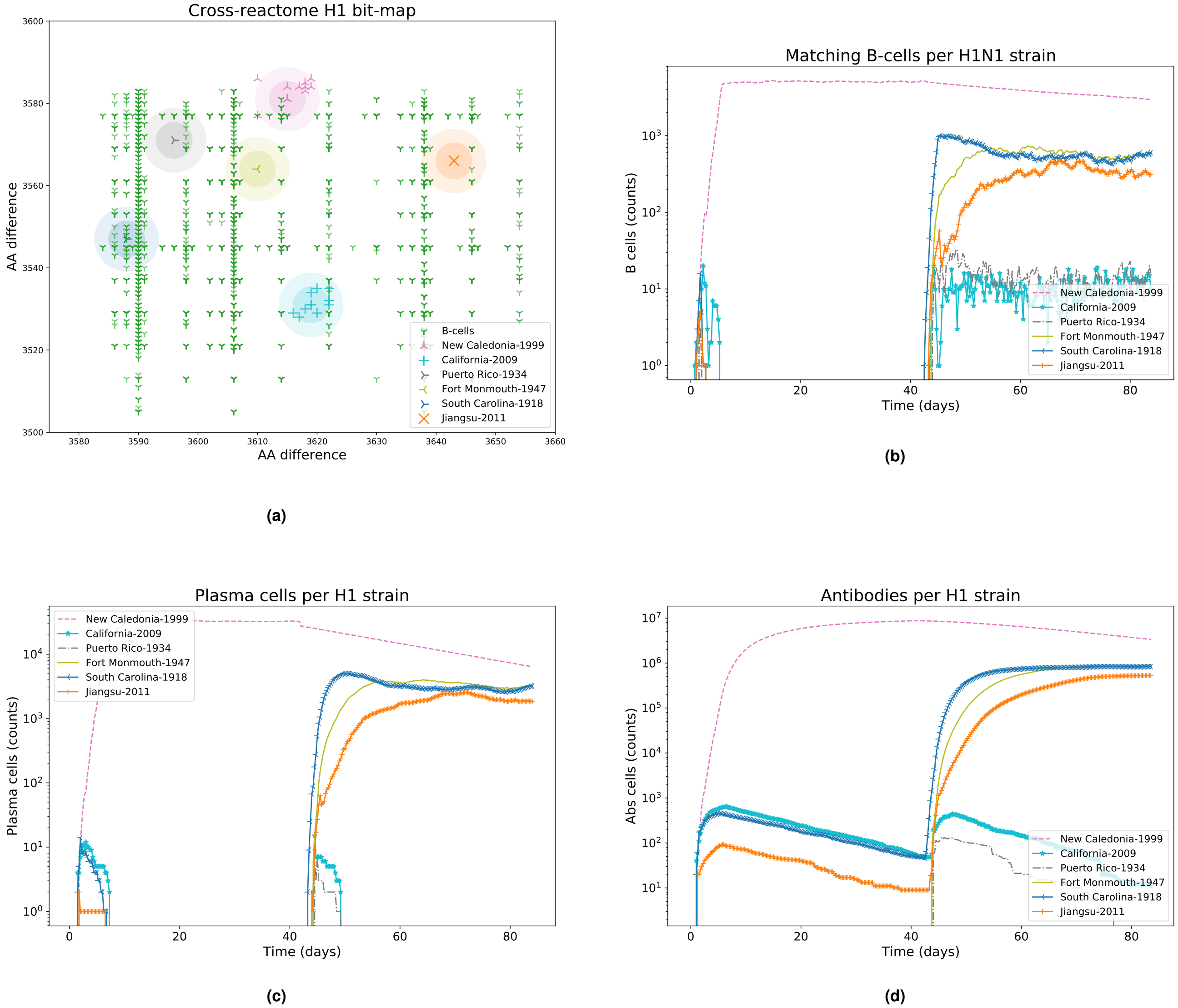


**Fig. (S32)** H1N1 Influenza strains *shape-space* and immune system components dynamics, Test 8.

Simulation 9 of the Principal Test with H1N1 strain.

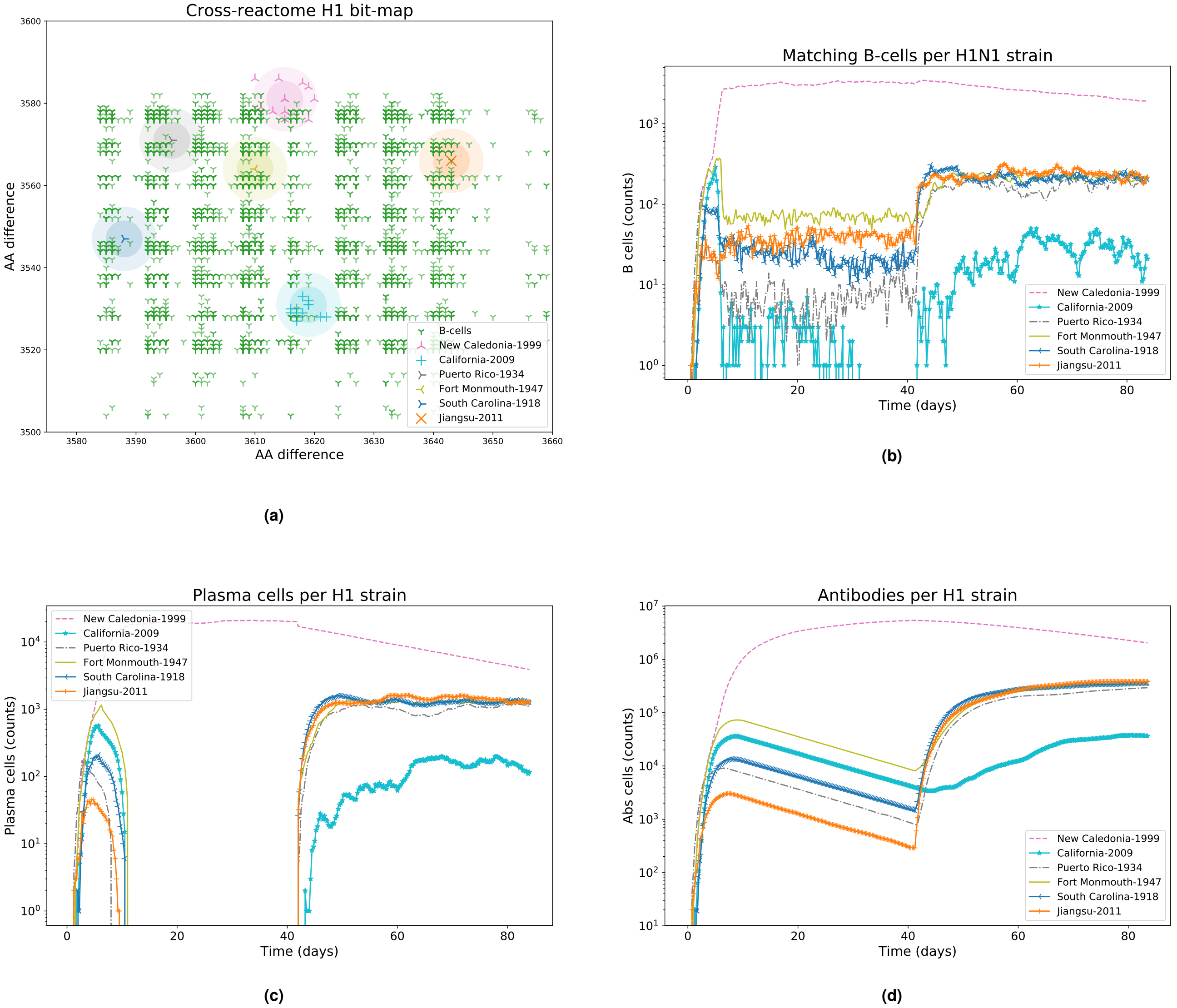


**Fig. (S33)** H1N1 Influenza strains *shape-space* and immune system components dynamics, Test 9.

Simulation 10 of the Principal Test with H1N1 strain.

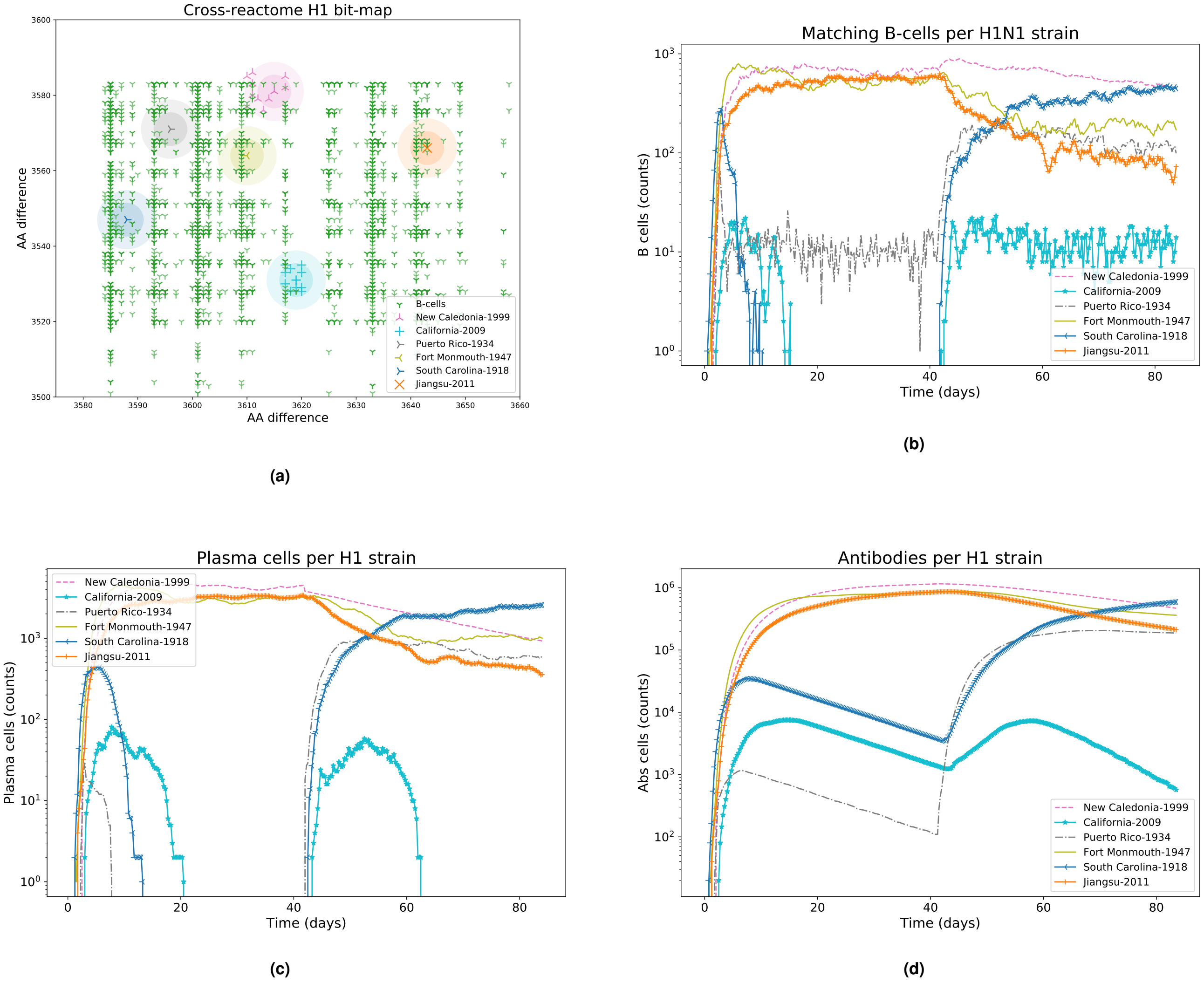


**Fig. (S34)** H1N1 Influenza strains *shape-space* and immune system components dynamics, Test 10.
